# Systematic analysis of transcriptional and epigenetic effects of genetic variation in Kupffer cells enables discrimination of cell intrinsic and environment-dependent mechanisms

**DOI:** 10.1101/2022.09.22.509046

**Authors:** Hunter Bennett, Ty D. Troutman, Enchen Zhou, Nathanael J. Spann, Verena M. Link, Jason S. Seidman, Christian K. Nickl, Yohei Abe, Mashito Sakai, Martina P. Pasillas, Justin M. Marlman, Carlos Guzman, Mojgan Hosseini, Bernd Schnabl, Christopher K. Glass

**Affiliations:** Department of Cellular and Molecular Medicine, University of California, San Diego, La Jolla, CA 92093, USA; Department of Medicine, University of California, San Diego, La Jolla, CA 92093, USA; Division of Allergy and Immunology, Center for Inflammation and Tolerance, Cincinnati Children’s Hospital Medical Center, University of Cincinnati College of Medicine, Cincinnati, OH 45229, USA; Metaorganism Immunity Section, Laboratory of Host Immunity and Microbiome, National Institute of Allergy and Infectious Diseases, NIH, Bethesda, MD 20892, USA; Department of Biochemistry and Molecular Biology, Nippon Medical School, 1-1-5 Sendagi, Bunkyo-ku, Tokyo 113-8602, Japan; Division of Allergy and Immunology, Cincinnati Children’s Hospital Medical Center, Cincinnati, OH 45229, USA; Department of Pathology, University of California, San Diego, CA, 920923 US; Department of Medicine, VA San Diego Healthcare System, San Diego, CA 92161, USA.

**Author notes:** These authors contributed equally and are ordered alphabetically (Hunter Bennett, Ty D. Troutman, Enchen Zhou).

## Abstract

Noncoding genetic variation is a major driver of phenotypic diversity but determining the underlying mechanisms and the cell types in which it acts remain challenging problems. Here, we investigate the impact of natural genetic variation provided by phenotypically diverse inbred strains of mice on gene expression and epigenetic landscapes of Kupffer cells. Analysis of gene expression in Kupffer cells and other liver cell types derived from C57BL/6J, BALB/cJ and A/J mice provided evidence for strain-specific differences in environmental factors influencing Kupffer cell phenotypes, including preferential Leptin signaling in BALB/cJ Kupffer cells. Systematic analysis of transcriptomic and epigenetic data from F1 hybrids of these mice, and transcriptomic data from strain-specific Kupffer cells engrafted into a common host enabled quantitative assessment of *cis* versus *trans* effects of genetic variation on gene expression and an estimate of cell autonomous versus non cell autonomous effects. Under homeostatic conditions, *trans* effects of genetic variation were dominant, with the majority of trans regulation being non cell autonomous. In contrast, strain specific responses to acutely administered LPS were primarily associated with genetic variation acting in *cis* to modify response elements for lineage determining and signal dependent transcription factors. Collectively, these findings reveal cell intrinsic and environmental effects of natural genetic variation on gene expression, demonstrate the use of enhancers as detectors of *trans* effects of genetic variation, and provide a new resource for understanding the impact of genetic variation on gene expression in Kupffer cells.

## INTRODUCTION

Genome-wide association studies have been highly successful in linking common forms of genetic variation to risk of disease and can provide important starting points for identification of new therapeutic targets. However, most of the genetic variants identified by these studies reside in non-coding regions of the genome, limiting their interpretability (Hindorff *et al.*, 2009; Farh *et al.*, 2015). Defining the causal variants, the cell types in which they exert their effects, and their mechanism of action remain major challenges. Common forms of non-coding genetic variation that include single nucleotide polymorphisms (SNPs) and short insertions and deletions (InDels) can alter gene expression by changing the sequences of DNA recognition elements for transcription factors within enhancers and promoters. For example, a SNP that reduces the binding of a required transcription factor to a cell-specific enhancer will result in reduced expression of the corresponding gene in that cell type. This represents a form of *cis*-regulation in which the variant exerts a cell- and gene-specific impact. Such variants can also have *trans* effects dependent on the gene that is affected. For example, a non-coding variant that affects the expression of a transcription factor in *cis* will also result in cell-autonomous *trans* effects on the target genes of that factor. Alternatively, if a non-coding variant affects expression of a gene that encodes a molecule involved in intercellular communication, such cell autonomous *cis* regulation can result in non-cell autonomous trans regulation in other cell types.

At the level of *cis* regulation, substantial progress has been made in linking SNPs and InDels associated with differential gene expression to promoters and cell-specific regulatory elements due to the increasing availability of cell-specific *cis*-regulatory atlases. Dependent on context, as much as 80% of allele-specific differences in *cis* regulatory activity can be linked to local variants. In contrast, there has been limited quantitative assessment of *trans* regulation in specific cell types of genetically diverse vertebrates in vivo (Veeken *et al.*, 2019; Veeken *et al.*, 2020; Zhong *et al.*, 2022). In addition, the systematic approaches for investigating mechanisms by which genetic variation exerts non-cell autonomous effects on cellular phenotypes are limited (Seldin *et al.*, 2018; Seldin, Yang and Lusis, 2019).

To address this gap, we integrated two distinct experimental strategies to distinguish cell autonomous and non-cell autonomous effects of genetic variation and infer altered signaling pathways associated with non-cell autonomous effects. The first strategy builds on the prior use of genetically diverse mice to investigate mechanisms of enhancer selection and activation. In these studies, macrophages were differentiated in vitro from bone marrow progenitor cells of five different inbred strains of mice and used for transcriptomic and genomic studies. Through an interrogation of the effects of SNPs and InDels on transcription factor binding, roles of approximately 80 transcription factors were inferred, including macrophage lineage-determining factors, signal-dependent transcription factors and other collaborative binding partners, in the selection and activation of macrophage-specific enhancers (Link *et al.*, 2018a). By establishing an identical differentiation program and cell culture environment, these studies largely excluded possible effects of genetic variation on other cell types and enabled a direct assessment of cell autonomous effects. In this context, studies in F1 hybrid mice indicated that up to 90% of strain specific enhancer activity exhibited a cis-pattern of regulation (Link *et al.*, 2018a; Hoeksema *et al.*, 2021; Veeken *et al.*, 2019).

The second experimental strategy is based on the roles of enhancers and promoters as sensors and transcriptional effectors of the internal and external signals that are necessary to establish cellular identity and function. Signal-dependent changes in gene expression generally result from altered binding and/or function of transcription factors at *cis* regulatory elements. The selection and activation of enhancers and promoters can be quantitatively measured on a genome-wide scale using assays for open chromatin and histone modifications associated with activity, such as acetylation of histone H3 at lysine 27 (H3K27ac) (Creyghton *et al.*, 2010). Motif enrichment analysis of enhancers and promoters that gain activity generally yields the sequences of binding sites for the transcription factors underlying gene activation. For example, quantitative analysis of dynamic enhancer landscapes in monocytes undergoing Kupffer cell differentiation in vivo revealed transcription factor motifs downstream of key signaling pathways in the liver that drive Kupffer cell specific gene expression, including Notch, TGFβ and LXR signaling pathways (Sakai *et al.*, 2019; Bonnardel *et al.*, 2019).

Here, we combine these experimental strategies to investigate mechanisms underlying cell autonomous and non-cell autonomous effects of naturally occurring genetic variation on enhancer activity and gene expression in Kupffer cells. Kupffer cells are the major population of resident macrophages in the liver and play important roles in various aspects of immunity and physiology, including detoxification of gut derived LPS and regulation of iron metabolism (Bennett *et al.*, 2021). Because they can be isolated in sufficient numbers to perform deep transcriptomic and epigenetic assays, they provide a powerful model system for evaluating the impact of natural genetic variation on tissue-resident macrophage populations *in vivo*. Kupffer cells are also implicated in numerous pathological processes involving the liver, including the development of non-alcoholic steatohepatitis (NASH) (Ramachandran *et al.*, 2019; Friedman *et al.*, 2018). To strengthen the resource value of these studies, we selected three strains of mice exhibiting different sensitivities to a NASH inducing diet for analysis of Kupffer cell gene expression (Hui *et al.*, 2015). We provide evidence that strain-specific differences in Kupffer cell transcriptomes and enhancer activity states can be used to infer consequences of genetic variation and their mechanism of action in other cell types that influence Kupffer cell gene expression. These findings suggest a general approach to investigation of non-cell autonomous effects of genetic variation that may be broadly applicable to diverse cell types.

## RESULTS

### Gene by environment interactions affecting Kupffer cell gene expression

To establish a model system for analysis of effects of natural genetic variation on Kupffer cell gene expression, we selected three common strains of inbred mice that recapitulate major phenotypic differences observed in human liver disease (Hui et al., 2015). Each chosen strain (A/J, BALB/cJ, and C57BL/6J) has a publicly available genome (Keane *et al.*, 2011) and positionally defined SNPs and Indels (Extended Data Fig. 1a). Comparisons of strain susceptibility/resistance to NASH confirmed documented trait segregation for development of obesity, steatosis, steatohepatitis, and fibrosis (Extended Data Fig. 1b-e). Transcriptional alteration of total hepatic Kupffer cells was most pronounced in NASH-susceptible strains, with minimal observed changes in cells from NASH-resistant mice (Extended Data Fig. 1f).

These findings prompted us to systematically evaluate the effects of genetic variation on the transcriptomes of Kupffer cells in these three strains of mice. In parallel, we generated transcriptomic data for bone marrow derived macrophages (BMDMs) from each strain to assess the extent to which effects of genetic variation were specific to Kupffer cells. Unsupervised clustering of this data demonstrated cell type as the dominant determinant of clustering, as expected, but each cell type also clearly segregated according to strain (Fig. 1a). Pairwise comparisons of Kupffer cells from each strain indicated between 194-362 differentially expressed genes (Fig. 1b-d). Comparing Kupffer cells to BMDMs, between 57 and 80 percent of genes with differential regulation between the selected strains were uniquely altered in Kupffer cells and not bone marrow derived macrophages or vice versa (Extended data Fig. 2a-c). Examples of transcripts with strain-unique patterns of expression in bone marrow macrophages are shown in Fig. 1e-g, and include notable regulators of inflammation (*Rgs1*, *Ifi203*, *Aoah*), responses to lipids (*Ch25h*, *Abcg1*, *Sdc1*), and polarization (*Arg2*, *Marco*). Likewise, examples for Kupffer cells are shown in Fig. 1h-j, and include transcriptional regulators (*Atf5*), suppressors of inflammation (*Cd300e*, *Irak3*), and drivers of inflammation (*Cxcl14*, *Cd40*, *Mefv*). Functional grouping of strain-unique gene expression by Kupffer cells linked evolutionary plasticity in programs controlling antigen processing and presentation, chemokine signaling, and suppression of chemotaxis (Fig. 1k) (Zhou *et al.*, 2019). These analyses provide insight into the functional effects of genetic diversification in regulating transcriptional programs by macrophages in two disparate environments.

**Fig. 1.**
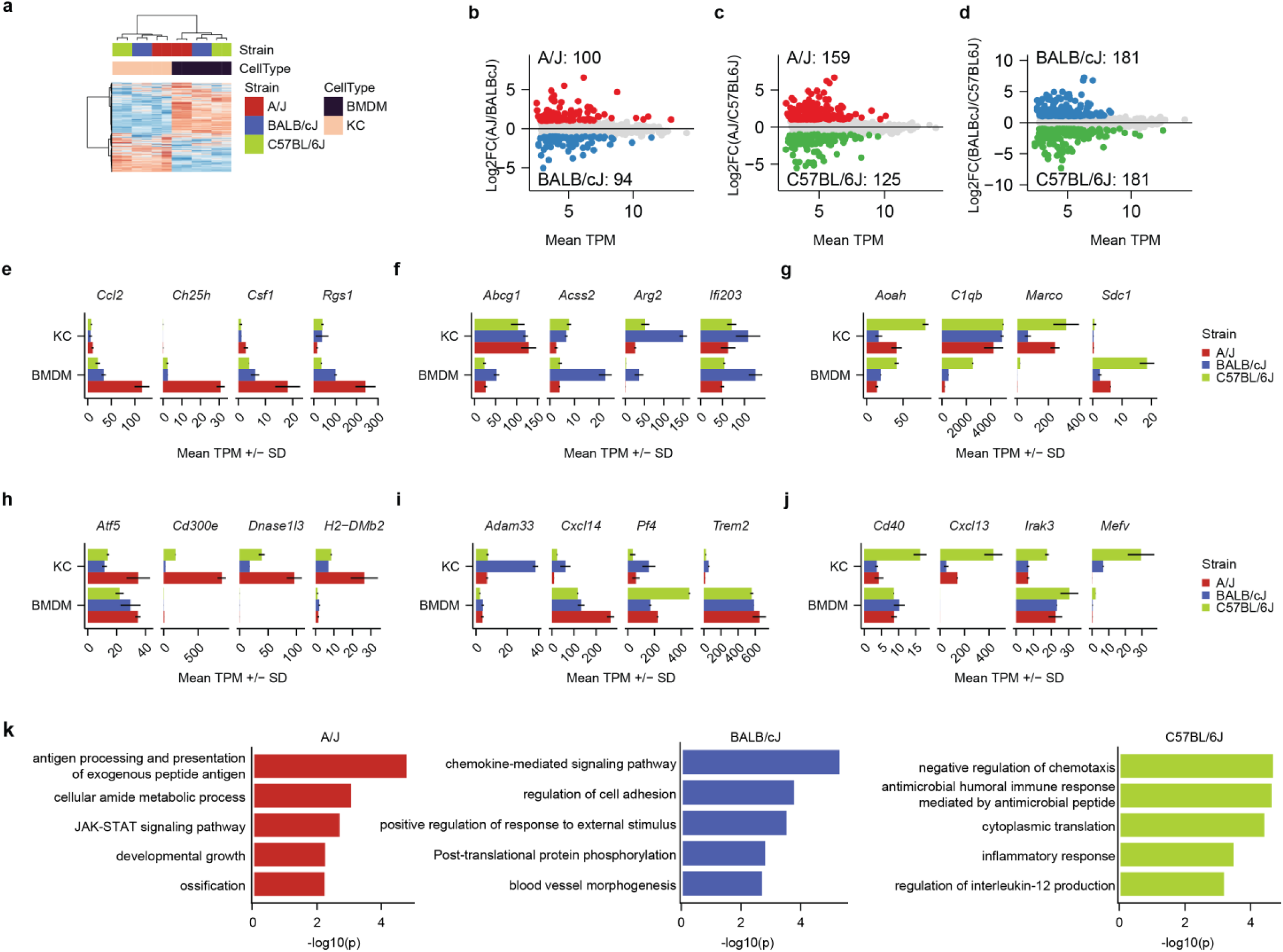
Gene-by-environment transcriptional regulation of primary mouse macrophages. **a**, hierarchical clustering of BMDM and Kupffer cell transcriptomes. Data represent any gene expressed above TPM threshold of 8. N=2 per group. **b-d,** comparison of the mean TPM and the DeSeq2 Log2FC for RNA-seq data from Kupffer cells purified from the indicated strains. Differentially expressed genes were identified using Log2FC > 1, adjusted p-value < 0.05, and TPM > 8 as identified using DeSeq2 (Log2FC and adjusted p-value) and HOMER (TPM normalization). **e-j,** expression of representative genes in BMDMs (**e, f, g**) and Kupffer cells (**h, i, j**) identified as A/J specific (left, **e, h**), BALB/cJ specific (middle, **f, i**), or C57BL/6J specific (right, **g, j**). Data represent the mean TPM +/- S.D. **k,** gene ontology enrichment for strain specific genes, which were defined as genes with significantly increased expression (Log2FC > 1, adjusted p-value < 0.05) in one strain compared to both other strains, e.g., increased expression in A/J relative to BALB/cJ and in A/J relative to C57BL/6J.

### Effect of natural genetic variation on the Kupffer cell enhancer landscape

To investigate the impact of genetic variation on potential regulatory elements in Kupffer cells, we performed assay for transposase accessible chromatin followed by sequencing (ATAC-seq) to identify regions of open chromatin that represent binding sites for transcription factors (Buenrostro *et al.*, 2013). Pairwise comparisons of ATAC-seq signal for each of the three strains are depicted in Fig. 2a. From ∼1000 to >7000 differential ATAC-seq peaks were identified, with the number of differential peaks scaling with the level of genetic diversity. Because the presence of open chromatin does necessarily reflect the activity of a putative regulatory element, we performed chromatin immunoprecipitation followed by sequencing (ChIP-seq) for H3K27ac, which is highly correlated with enhancer and promoter activity (Creyghton *et al.*, 2010). ATAC-seq peaks were then annotated with normalized H3K27ac tags using a 2000 bp window centered on ATAC-seq peaks. ATAC-peaks with at least log2>4 tags in at least one strain were considered potential transcriptional regulatory elements. Approximately 5000 to 7000 ATAC-seq peaks were found to have differential H3K27Ac signal per strain-by-strain comparison (Fig. 2b). More than 10,000 of the ∼66,000 putative regulatory elements identified were defined to be under genetic control. Venn diagrams of the overlaps of ATAC and H3K27ac annotated ATAC-seq peaks are provided in Fig. 2c. Examples of strain-specific ATAC-seq and H3K27ac ChIP-seq signals that correlate with strain specific gene expression are provided by *Cd300e*, *Trem2* and *Irak3* (Fig. 2d).

To estimate the extent to which local genetic variants contribute to strain-specific differences in ATAC-seq and H3K27ac, we determined the frequency of SNPs and InDels within ATAC-defined open chromatin at strain similar ATAC and H3K27ac peaks in comparison to peaks exhibiting more than a 2-fold change in ATAC or H3K27ac signal. Strain similar peaks exhibited a background frequency of SNPs and InDels of 15-18%, whereas the variant frequency was 46-56% in peaks differing by more than 2-fold (Fig. 2e). The variant frequency increased to 57-64% at a 4-fold difference between strains and was similar at an 8-fold difference. Overall, ∼30-50% of regions quantitative differences in open chromatin or histone acetylation were associated with nearby variants controlling chromatin opening or activity in *cis*.

Motif enrichment analysis of the common set of ATAC peaks exhibiting H3K27ac yielded motifs corresponding to previously established Kupffer cell lineage determining factors, including PU.1, MAF/MAFB, NFkB, TFEB/TFEC, LXR, RBPJ and SMADs (Extended Data Fig 3a-b). To identify potential transcription factors driving strain specific enhancer selection, we performed motif enrichment of strain specific enhancers. In each case, binding sites for PU.1 were the most enriched motifs, consistent with the general role of PU.1 in the selection of macrophage-specific enhancers (Fig. 2f). In addition, putative enhancers exhibiting preferential H3K27ac in A/J mice were enriched for motifs recognized by Nfil3 and CTCF, whereas corresponding enhancers in BALB/cJ Kupffer cells were enriched for AP-1, MAF, ATF3 and MITF motifs, and for Yy1, Yy2, Egr1 and Ets motifs in C57Bl/6J Kupffer cells (Fig. 2f). As this analysis considers all the preferentially active enhancers, a causal relationship between enrichment of a motif and enhanced enhancer activity would most likely be due to increased activity of corresponding transcription factors in that strain.

To gain further insight into mechanisms by which SNPs and InDels exert local effects on enhancer selection and function, we assessed the quantitative impact of the genetic variation provided by these three strains of mice on the open chromatin and H3K27ac using the motif mutation analysis tool (Shen *et al.*, 2020). MAGGIE associates changes of epigenomic features at homologous sequences (e.g., enhancer activation or enhancer repression) with motif mutations caused by genetic variation to prioritize motifs that likely contribute to the local regulatory function. Although the extent of genetic variation provided by the comparison of A/J, BALB/cJ and C57BL/6J is substantially less than that used in previous applications of MAGGIE (Hoeksema *et al.*, 2021; Shen *et al.*, 2020), systematic analysis of JASPAR (Mathelier *et al.*, 2016) transcription factor binding motifs identified more than 100 motifs for which variants were significantly associated with gain or loss of ATAC-seq and/or H3K27ac signal (**Supplemental Table 1**). Many motifs identified by MAGGIE are binding sites for related transcription factors, e.g., Ets, AP-1 and IRF families. A representative selection of these motifs is presented in Fig. 2g. Overall, effects of motif mutations on open chromatin and H3K27ac were highly correlated. Significance of motif mutation enrichments associated with ATAC-seq peaks differences were greater that what we detected with H3K27ac differences, likely due to sample size differences between these subsets. Mutations affecting PU.1 and related Ets family motifs were the most deleterious for chromatin accessibility and acetylation, consistent with role of PU.1 as a macrophage lineage determining transcription factor (Heinz *et al.*, 2010; Link *et al.*, 2018a; DeKoter and Singh, 2000; Scott *et al.*, 1994). Motif mutations affecting IRF, STAT, AP-1/ATF, CREB, MAF, and C/EBP families were also identified as drivers of enhancer selection and function in Kupffer cells. The identification of a motif for LXR/RXR heterodimers is also consistent with its established role as a Kupffer cell lineage determining factor that participates in chromatin opening and enhancer activation (Sakai et al., 2019). Not all enriched motif mutations could be associated with corresponding transcription factors expressed by Kupffer cells (e.g. ZFP57, ZKSCAN, PAX and NKX, denoted by #), suggesting their involvement in other biological contexts, or the interaction of transcription factors as yet undefined to interact with these DNA elements. Rbpj and SMAD both regulate Kupffer cell differentiation (Sakai *et al.*, 2019; Bonnardel *et al.*, 2019), however the abundance of SNPs and InDels affecting these motifs was limited. Consequently, MAGGIE analysis did not detect enrichment of Rbpj or SMAD affecting motif mutations associated with altered chromatin accessibility or acetylation. This finding may reflect a selection pressure to preserve the function of these elements. In concert, these studies establish the impact of a moderate level of natural genetic variation on the enhancer landscapes of Kupffer cells and provide support for functional roles of the major motifs found to be enriched in these regulatory elements.

**Fig. 2.**
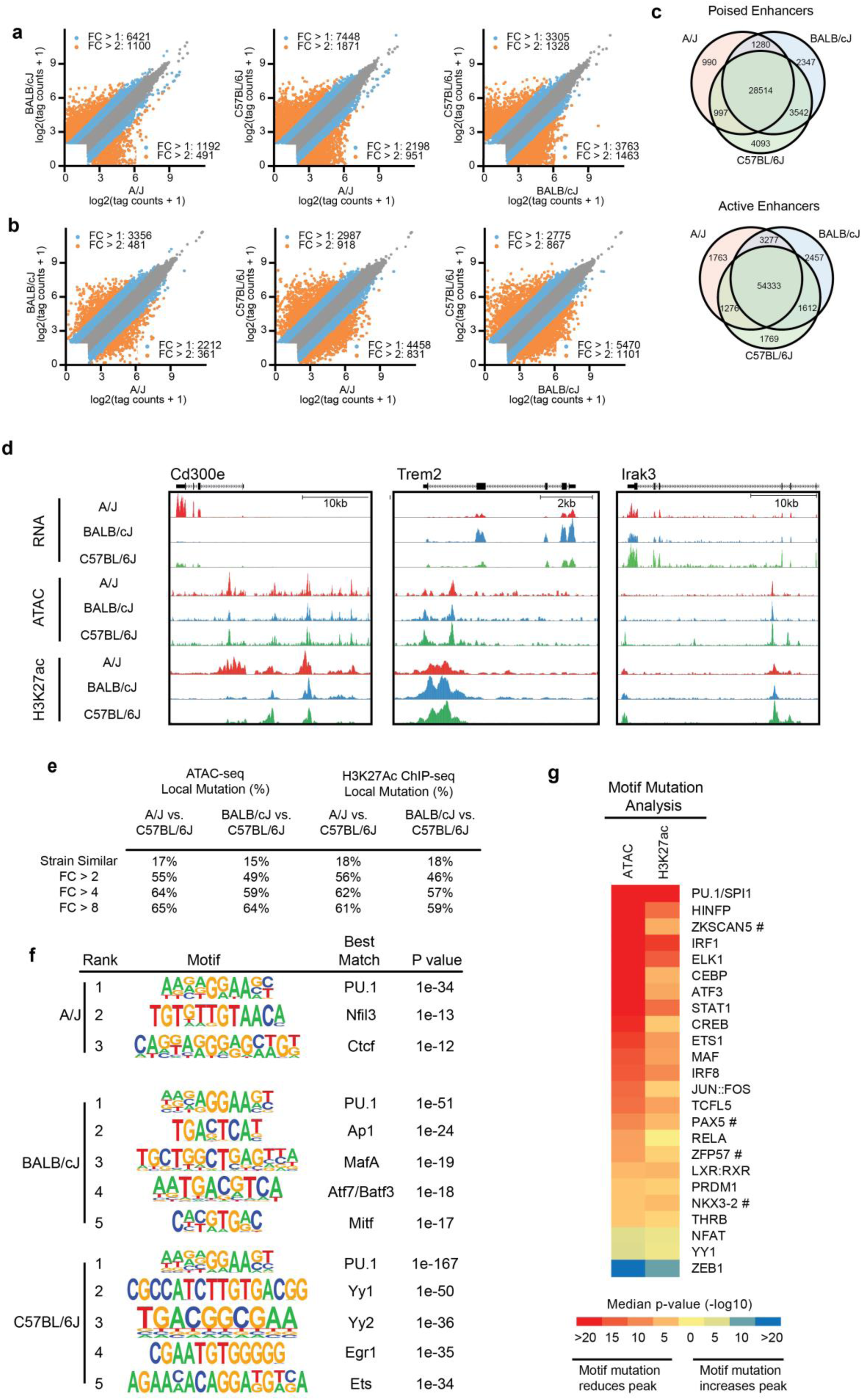
Local and global effect of natural genetic variation on Kupffer cell epigenome. **a**, scatterplots of log2 tag counts for ATAC-seq signal at union set of IDR ATAC-seq peaks across all strains. **b,** scatterplots of log2 tag counts for H3K27Ac ChIP-seq signal at union set of IDR ATAC-seq peaks across all strains, tags annotated with a window size of 2000bp centered on middle of IDR peak. **c**, overlap of active and poised genomic loci for each strain. Poised loci defined as sites with >8 HOMER normalized ATAC-seq tags. Active loci defined as sites with >32 HOMER normalized H3K27Ac ChIP-seq tags. **d,** strain specific epigenetic signals associated with transcriptional activation. *Cd300e* expression and H3K27Ac acetylation of nearby enhancers are specific to A/J Kupffer cells. *Trem2* is preferentially expressed in BALB/cJ Kupffer cells and associated with increased acetylation of an intronic enhancer. *Irak3* is preferentially expressed in C57BL/6J Kupffer cells and is associated with C57BL/6J specific ATAC-seq peaks and increased acetylation of nearby enhancers. **e,** enhancers were categorized into strain-similar or strain-specific for both ATAC-seq and H3K27Ac ChIP-seq data. Table denotes percentage of enhancers at each fold change cutoff that harbor local genetic variation within the 200bp IDR peak. **f,** top motifs associated with strain specific active enhancers, defined as loci that had strain specific increases in H3K27Ac. **g**, MAGGIE motif mutation analysis on strain differential open and active enhancers.

### Inference of environmental signals influencing strain-specific Kupffer cell gene expression

Sinusoidal endothelial cell expression of *Dll4* acting on the Notch pathway and stellate cell expression of *Bmp9* acting on the ALK/SMAD pathway provide signals required to drive the expression and activity of Kupffer cell lineage determining factors (Sakai *et al.*, 2019; Bonnardel *et al.*, 2019; Guilliams *et al.*, 2022). To gain insights into how these and other signaling molecules affect strain-specific transcription in Kupffer cells we applied the NicheNet, a computational model of intercellular signaling (Browaeys, Saelens and Saeys, 2020). We performed RNA-seq on hepatocytes, stellate cells, and liver sinusoidal endothelial cells (LSECs) from A/J, BALB/cJ, and C57BL6/J mice and asked whether ligands expressed by could predict strain specific Kupffer cell gene expression patterns (Fig. 3a, Extended Data Fig. 4). We also included a selection of hormones that could potentially alter transcription because Kupffer cells are readily exposed to portal blood.

The top scoring NicheNet ligands are summarized in Fig. 3b and a circos plot summarizing ligand-target gene connections is shown in Fig. 3c. Hepatocyte derived ligands included *ApoE*, an apolipoprotein that binds lipoprotein receptors including the low-density lipoprotein receptor (Ldlr) and Trem2 and was predicted to induce BALB/cJ specific Kupffer cell gene expression of genes including *Fads1* and *Cxcr4* (Fig. 3c). LSEC derived ligands included *Bmp2*, a member of the BMP ligand family, and predicted to expression of genes specific to A/J Kupffer cells. *Adam17* and *App* were identified as niche ligands provided by LSEC and hepatic stellate cells. *App*, or amyloid precursor protein, predicted C57BL/6J specific gene expression, including inflammation response genes *Ccl5* an*d Tnfaip3*. *Lep*, encoding the adipokine Leptin, was one of the top scoring ligands predicting strain-specific gene expression in BALB/cJ Kupffer cells, and *Lepr,* which encodes the leptin receptor, was expressed highest in BALB/cJ Kupffer cells (Fig 3d). Altogether, these results predicted alteration in expression or activity of several niche ligand:receptor signaling pathways regulating strain-specific Kupffer cell gene expression.

To validate one of these predictions, we investigated the significance of differential expression of *Lepr*, which is expressed highest by Kupffer cells in the hepatic niche (Fig. 3d). Leptin binding to the Leptin receptor results in phosphorylation and activation of STAT3 (Imajo et al., 2012; Kiguchi et al., 2009; Maeda et al., 2009) We found that acute intraperitoneal injection of leptin into fasted mice induced detectable STAT3 phosphorylation in the livers of C57BL/6J and BALB/cJ mice. However, A/J mice, which had the lowest *Lepr* expression by Kupffer cells (Fig. 3d), had undetectable STAT3 phosphorylation (Fig. 3e). This result is consistent with the prediction of Niche-Net that leptin signaling could contribute to strain specific differences in Kupffer cell gene expression. Prior studies have shown that the leptin-STAT3 pathway can alter inflammatory signaling in obese mice (Imajo et al., 2012) and that Kupffer cells facilitate the acute effects of leptin on hepatic lipid metabolism (Metlakunta *et al.*, 2017). Analysis of *Lepr* expression in C57BL/6J mice in the context of a NASH-inducing diet indicate that it is strongly downregulated in resident (Tim4+) Kupffer cells and is not detectable in blood monocytes or recruited monocyte derived (Tim4-) Kupffer cells that accumulate during the progression of NASH (Fig. 3f). *Lepr* is also one of the few genes that are not induced in monocyte-derived cells that repopulate the Kupffer cell niche following experimental ablation of resident Kupffer cells (Fig. 3g). Thus, *Lepr* expression in Kupffer cells is strongly determined by both developmental origin and environment.

**Fig. 3.**
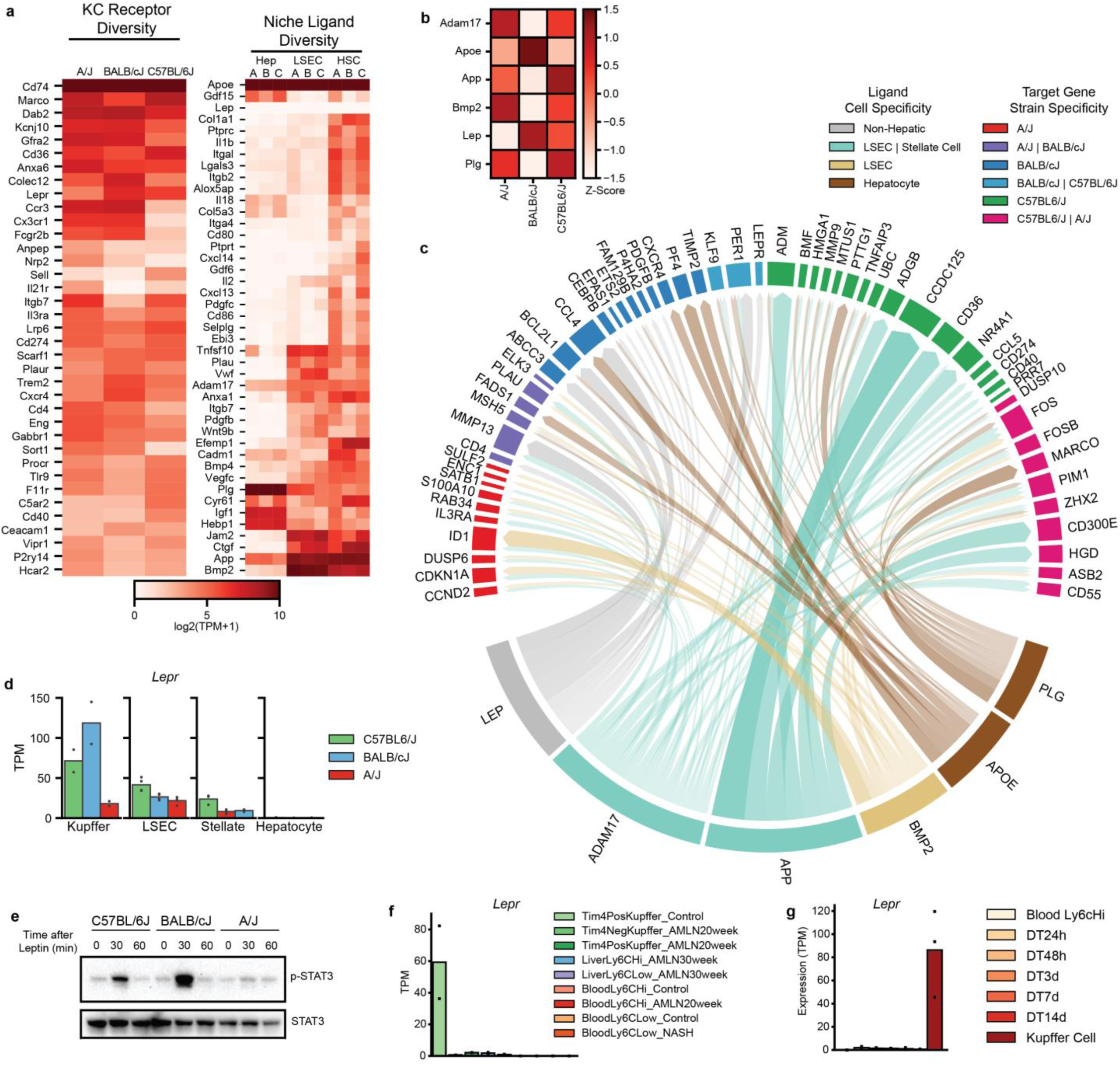
Hepatic Kupffer cell-niche differences predicted using network analysis. **a**, gene expression of receptors by Kupffer cells (left) or ligand by niche companion cells (right) from the indicated cell types (A, B, C in righthand plot indicate expression in A/J, BALB/cJ, and C57BL/6J, respectively). **b,** top 2 NicheNet ligand activity scores for each strain, significance normalized to z-score across strains. **c**, circos plot demonstrating gene targets of top 6 NicheNet ligands from b. **d**, strain specific expression of the leptin receptor in hepatic cells. **e**, immunoblot assessment of total and phosphorylated STAT3 in liver tissue from mice injected with 1mg/kg leptin via intraperitoneal route. Data are representative of 3 experiments. **f,** expression of leptin receptor in Kupffer cells from healthy C57BL/6J mice (left) and myeloid cells including macrophages and monocytes isolated from mice fed an AMLN NASH-inducing diet for 20 weeks. **g**, expression of leptin receptor in embryonic-derived Kupffer cells (far right), and bone marrow derived monocytes repopulating liver at specific time points following depletion of resident Kupffer cells with diptheria toxin.

### Cis and trans effects of genetic variation on Kupffer cell gene expression

We next used RNA-seq data from two complementary model systems to further clarify the mechanisms by which genetic variation provided by C57BL/6J and BALB/cJ mice exerted effects on gene expression. First, we transplanted bone marrow from C57BL/6J mice or BALB/cJ mice into busulfan conditioned NOD-Scid-gamma (NSG) recipients (Fig. 4a) (Peake *et al.*, 2015). Busulfan treatment also depletes resident Kupffer cells, allowing monocyte-derived Kupffer cells from donor progenitors. In this model, donor-derived Kupffer cells from chimeric NSG mice reside within matched host environments but retain their strain specific genomes. The second model used mice from the first-generation intercross of C57BL/6J male mice and BALB/cJ female mice (CB6F1/J mice, denoted as F1-hybrid) (Fig. 4a). In Kupffer cells from F1-hybrid mice, the autosomal parental BALB/cJ and C57BL/6J genomes co-exist within a matched extracellular and intracellular environment. To discern allelic bias from F1-hybrid RNA-seq data we mapped the raw sequence to each parental genome and compared the levels of perfectly mapped reads spanning mutations between the parental strains, as described previously (Heinz *et al.*, 2013; Link *et al.*, 2018a; Link *et al.*, 2018b; Hoeksema *et al.*, 2021).

In this experimental design, F0 strain-specific gene expression that is retained in NSG Kupffer cells and in an allele-specific manner in F1-hybrid Kupffer cells is interpreted to result from *cis* acting genetic variation. Strain-specific expression that is lost in NSG Kupffer cells is interpreted to result from non cell-autonomous *trans*-acting genetic variation, i.e., through effects of genetic variation on factors that are extrinsic to Kupffer cells and that differ between the C57BL/6J and BALB/cJ strains. Conversely, strain specific gene expression that is retained in the NSG Kupffer cells but lost in the F1-hybrid Kupffer cells is interpreted to result from cell-autonomous or non cell-autonomous *trans*-acting genetic variation.

Gene expression differences are illustrated for each model in Fig. 4b and Fig. 4c. We identified differential expression of ∼240 genes between engrafted Kupffer cells of each strain. Next, we assessed if residence of Kupffer cells from discrete genetic backgrounds in a matched environment leads to transcriptome convergence by intersecting differential expression lists between Kupffer cells from F0 BALB/cJ or C57BL/6J Kupffer cells (Fig. 1d) and NSG chimeras (Fig. 4b). This analysis revealed that 58% (211/361) of differentially expressed gene between Kupffer cells from F0 BALB/cJ or C57BL/6J mice were not differentially expressed in livers of NSG chimeras, while 150 genes had strain-specific expression patterns in both the parental mice and the NSG model. One explanation could be that engrafted cells are of hematopoietic origin in contrast to embryonic origin expected of Kupffer cells in from healthy parental mice (Scott *et al.*, 2016). We compared the set of strain specific genes in F0 mice but lost in NSG to the set of genes that are differentially expressed between embryonic Kupffer cells and hematopoietic Kupffer cells using previously published data (Sakai *et al.*, 2019). Of the 211 NSG *trans* genes, only 19 displayed decreased expression in hematopoietic Kupffer cells compared to embryonic Kupffer cells. We therefore interpret these results to indicate that differential expression of genes in Kupffer cells from parental strains is driven primarily by non-cell autonomous factors differing between C57BL/6J and BALB/cJ mice.

Kupffer cells from the NSG and F1-hybrid mice displayed similar levels of strain bias (245 DEGs) using identical statistical thresholds (Fig. 4b, c). This was surprising because only 4407 genes could be confidently mapped with allelic bias due to the limited genetic diversity between transcripts from each genome, whereas in the engraftment model, all 8420 transcripts that met a minimum TPM threshold of 8 were considered. However, in the F1-hybrid model, ∼75% (278/361) of genes expressed differentially by Kupffer cells from parental strains were detected at similar levels between alleles from F1-hybrid cells. To more accurately quantify the relative contribution of *cis*, cell autonomous *trans*, and cell-non autonomous *trans* variation, we compared the fold change strain-specific genes between parental Kupffer cells to cells from the NSG and F1-hybrid models (Fig. 4d, e). We defined *cis* regulated genes to be genes that displayed significant (DeSeq2: log2 fold change > 1, adjusted p value < 0.05) strain specific bias in the same direction, *trans* genes as genes that displayed strain specific bias in the parental cells but not the NSG or F1-hybrid models, and *mixed* genes as those that displayed strain specific bias in the NSG or F1-hybrid models but not parental cells. We found 172 *trans*-regulated genes in NSG chimeras, and 124 *trans*-regulated genes in the F1-hybrid Kupffer cells (Fig. 4d, e).

We excluded genes lacking mutations to consider genes with detectable allelic bias in F1-hybrid cells (Fig. 4f) and intersected the resulting gene lists. We found that the majority (79/125) of transcripts with established allelic bias were *trans*-regulated in both models and likely driven by environmental *trans* effects in the parental hepatic environment that are lost in both the NSG and F1-hybrid models. Many remaining transcripts (38/125) were *trans*-regulated in F1-hybrids but not the engraftment model, suggesting they are caused by cell-autonomous differences upstream of transcription factor binding. A small proportion (8/125) genes were *trans*-regulated in NSG chimeras but not F1-hybrids. Examples of each type of *trans* gene are shown in Fig. 4g. Based on these data, we estimate that 2/3 of *trans* mediated genetic variation in Kupffer cells is driven by cell non-autonomous differences in environmental signals, while 1/3 is due to cell-autonomous differences in signaling or transcriptional activity. Notably, Gene ontology analysis indicated that trans genes specific to C57BL/6J Kupffer cells were functionally enriched annotations involved in antigen presentation and response to lipopolysaccharide, including *Aoah, Lbp, Cd40, Batf3, Cd74* and *Tnfaip3*, whereas trans genes specific to BALB/cJ Kupffer cells were enriched for cell removal and chemokine signaling programs, including *Cd209f, Cd209g*, *Cxcr4* and *Pf4* (Fig. 4i).

**Fig. 4.**
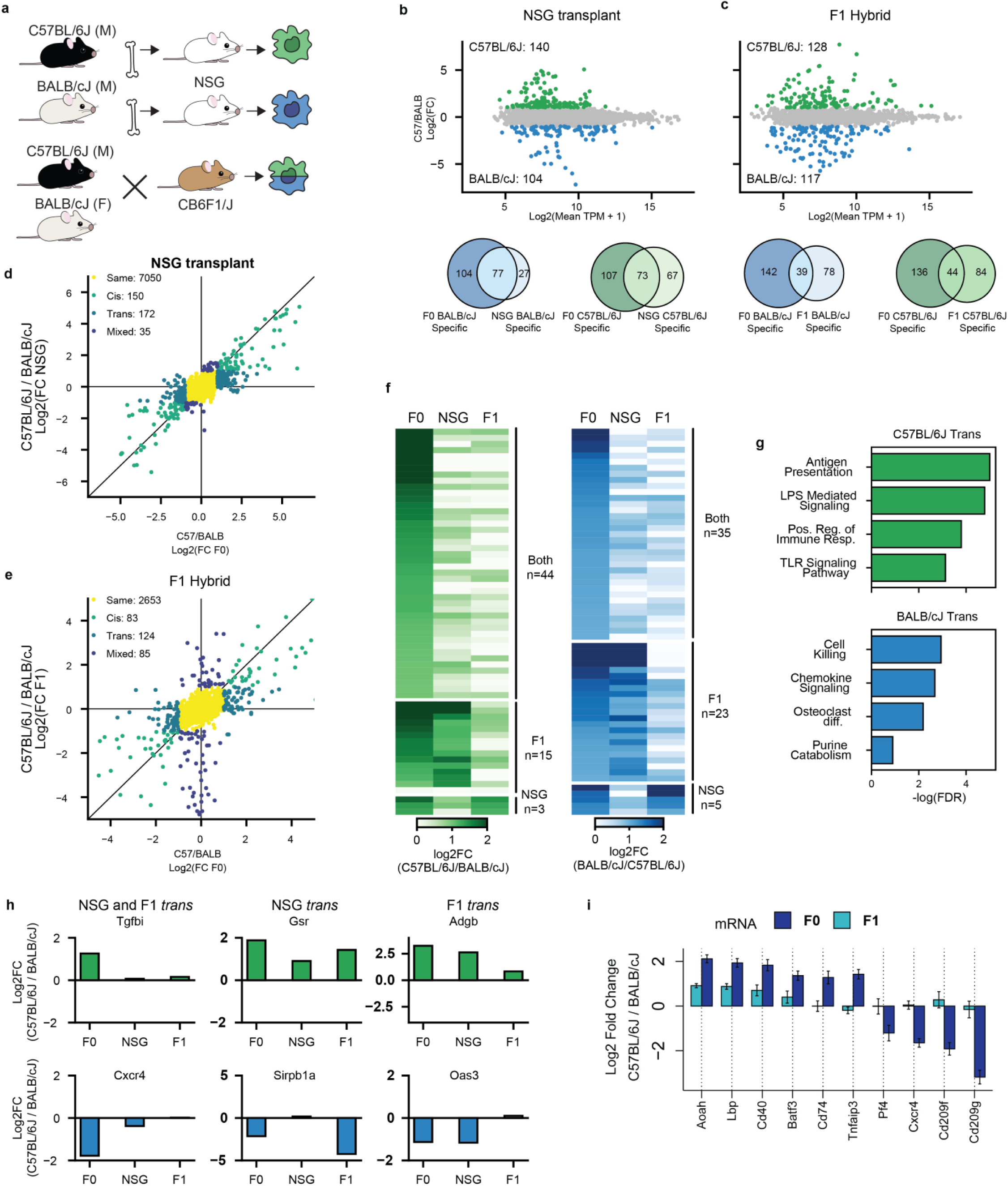
Cell autonomous and non-autonomous *trans* interactions contribute to gene expression differences in Kupffer cells. **a**, experimental schematic for chosen strategies used to predict cell autonomous and non-cell autonomous patterns of strain specific gene expression by Kupffer cells. (Top) Immune deficient NOD/Scid-IL2rg (NSG) mice were treated with busulfan and transplanted with hematopoietic stem cells of either strain as indicated. Donor-derived Kupffer cells from engrafted mice were isolated using FACS based on HLA haplotype. (Bottom) F1-hybrids resulting from the intercross of female BALB/cJ mice with male C57BL/6J mice. Kupffer cells from this hybrid strain, referred to as CB6F1/J, were purified by FACS. **b,** differential gene expression between BALB/cJ and C57BL/6J Kupffer cells isolated from NSG mice. Data are presented in an ‘MA plot’ format. On the x-axes, data depict the log2 transformed average TPM between samples from the displayed comparison. On the y-axes, data depict relative expression differences (Log2 fold change) calculated using DeSeq2. Venn diagram shows degree of overlap between differential BALB/cJ or C57BL/6J specific genes from F0 Kupffer cells and NSG transplanted Kupffer cells. **c,** Comparison of BALB/cJ and C57BL/6J allelic mRNA expression in CB6F1/J F1 mice, data displayed as described in b. **d-e**, comparison of gene expression ratios using Kupffer cells from parental mice (x-axes) to Kupffer cells from transplant recipients (**d**) or F1-hybrids (**e**). Genes with maintained expression differences in both comparisons are labeled as ‘Cis’ and colored green; genes differentially expressed in parental cells but not in the indicated environmental model are labeled as ‘Trans’ and colored blue; genes expressed similarly in parental data and differentially in the environmental models are labeled as ‘Mixed’ and colored purple; genes expressed similarly in both comparisons are labeled ‘Same’ and colored yellow. **f**, overlap of *trans* genes identified in F1 and NSG models. *Trans* genes were identified as in d and e. NSG *trans* genes were filtered to only consider genes harboring a mutation allowing discrimination of allelic reads in F1 Kupffer cells. *trans* genes were sorted into genes shared with F1 and NSG and genes that were *trans* only in F1 or only in NSG. **g**, metascape enrichment of C57BL/6J and BALB/cJ specific *trans* genes from F1 to F0 comparison. **h**, log2 transformed fold change of genes within each of the categories of *trans* gene described in g. **i**, Log2 transformed fold change values for selected C57BL/6J and BALB/cJ specific *trans* genes.

### Cis and trans effects of natural genetic variation on Kupffer cell enhancers at the steady state

These findings are consistent with genetically determined differences in the hepatic environment exerting an additional layer of regulation on Kupffer cells *in vivo*. Our results here predicted that *trans* effects of genetic variation would be dominant drivers of strain specific differences in open chromatin and histone acetylation in Kupffer cells. To test this hypothesis, we performed ATAC-seq and ChIP-seq for H3K27ac in Kupffer cells isolated from CB6F1/J F1-hybrid mice and evaluated data with allelic bias as discriminated by the presence of genetic variants. As with gene expression, *trans* effects were dominant for both ATAC-seq peaks (442 cis and 1184 trans, Fig. 5a) and overlayed H3K27ac ChIP-seq signal (936 cis and 1493 trans, Fig. 5b) (DeSeq2: log2 fold change >1, adjusted p-value < 0.05) (Love, Huber and Anders, 2014). Representative loci exhibiting *cis* regulation of open chromatin and H3K27ac are illustrated for *Plac8* and *Ifi202b* in Fig. 5c. Conversely, *trans* regulation of open chromatin and H3K27ac are exemplified in the vicinity of *Cd74* (Fig. 5c). Although changes in open chromatin and H3K27ac were generally highly correlated, the *Cxcr4* locus provides an example of strain-similar ATAC-seq signal but BALB/cJ-preferential H3K27ac signal that is equalized in the F1-hybrid Kupffer cells (Fig. 5c).

To identify transcription factors driving *trans* regulation of open chromatin and H3K27ac, we performed motif enrichment analysis of the *trans* regulated 1184 ATAC peaks and 1493 H3K27ac loci with convergent allelic signals in F1-hybrid Kupffer cells. Motifs for Ets factors were preferentially enriched in trans-regulated regions of open chromatin specific to BALB/cJ Kupffer cells, while motifs for CTCF and IRFs were preferentially enriched in open chromatin specific to C57BL/6J Kupffer cells (Extended data Fig. 5a). ATAC-seq peaks exhibiting higher levels of H3K27ac in BALB/cJ Kupffer cells that are strain-similar in the F1-hybrid had strong preferential enrichment for motifs recognized by AP-1 factors, Maf factors and Foxo1, whereas motifs recognized by IRF, LXR, and NFκB were enriched in C57BL/6J specific H3K27Ac regions that are strain-similar the F1-hybrid (Fig. 5d). LXRα is more highly expressed in C57BL/6J Kupffer cells (Fig. 5e), potentially explaining enrichment of the LXRE in these strain specific chromatin regions. However, the majority of transcription factors associated with strain specific ATAC or H3K27ac peaks were similarly expressed between strains (Fig. 5e). These observations implicate signaling pathways upstream of AP-1 and Maf factors as more active in BABL/cJ Kupffer cells, and pathways upstream of NFκB as more active in C57BL/6J Kupffer cells.

**Figure 5.**
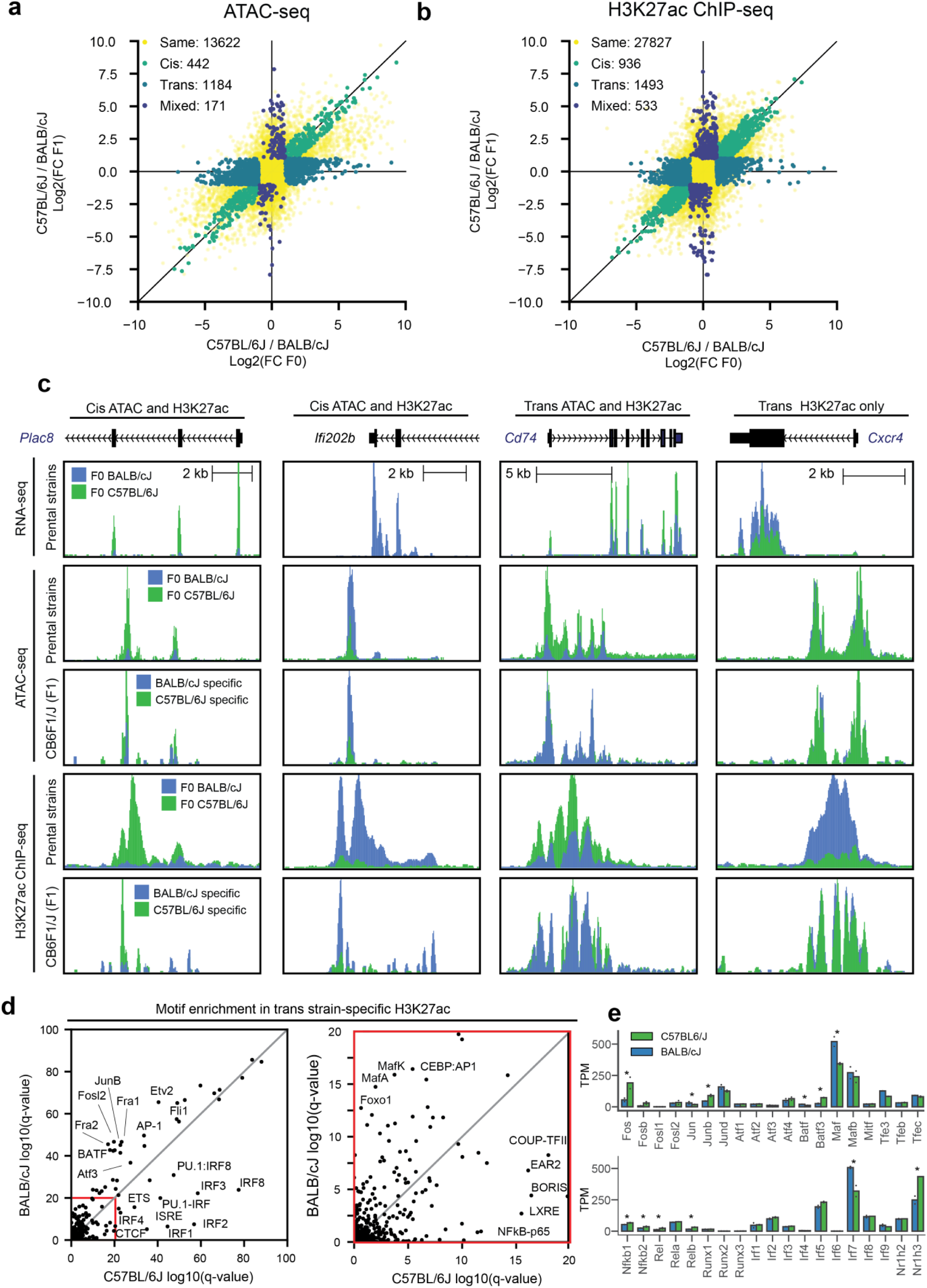
*Cis* and *trans* analysis of epigenetic loci reveals upstream regulators of *trans* transcriptional diversity. **a**, comparison of ATAC-seq tags ratios using Kupffer cells from parental mice (x-axes) and F1-hybrids (y-axis). All IDR ATAC-seq peaks harboring a mutation were considered for analysis. Peaks with maintained ATAC-seq tag differences in both comparisons are labeled as ‘Cis’ and colored green; peaks with differential ATAC-seq tags in parental cells but not in the F1 model are labeled as ‘Trans’ and colored blue; peaks with similar ATAC-seq tags in parental data and differential ATAC-seq tags in the F1 are labeled as ‘Mixed’ and colored purple; peaks with similar ATAC-seq tags in both comparisons are labeled ‘Same’ and colored yellow. **b**, comparison of H3K27Ac ChIP-seq tag ratios using Kupffer cells from parental mice and F1 hybrids using methods described in **a**. **c**, examples of *cis* and *trans* epigenetic loci. **d**, enrichment score of known transcription factor motifs in *trans* BALB/cJ specific enhancers (y axis) and in *trans* C57BL/6J H3K27Ac enhancers (x axis). **e**, expression of selected transcription factors in C57BL/6J and BALB/cJ F0 Kupffer cells. * indicates differential expression with FDR adjusted p-value < 0.05.

### Impact of genetic variation on Kupffer cell responses to LPS

The different sensitivities of C57BL/6J and BALB/cJ mice to a NASH-inducing diet (Extended Data Fig. 1) implies strain specific differences in cell type responses within and outside of the liver during the dietary intervention. In the context of NASH models, the effects of diet on Kupffer cell gene expression may be indirect, as substantial changes in gene expression in C57BL/6J Kupffer cells may not occur until after development of histological changes in the liver (Santhekadur, Kumar and Sanyal, 2018). To gain insights into how genetic variation affects Kupffer cell responses to an environmental perturbation *in vivo*, we assessed changes in the transcriptomes and open chromatin landscapes Kupffer cells from BALB/cJ and C57BL/6J mice two hours following intraperitoneal injection of lipopolysaccharide (LPS), a time point capturing immediate transcriptional consequences of TLR signaling. We found a total of 3268 genes were differentially regulated by LPS injection in Kupffer cells from the two parental strains (DeSeq2: log2 fold change >1, adjusted p-value < 0.05, minimum average TPM > 8). Comparison of log2 fold changes indicated a conserved response of 1885 genes, and strain divergence in 1383 DEGs responsive in only one strain (Fig. 6a). We also observed substantial LPS responsive changes in chromatin accessibility (DeSeq2: log2 fold change >1, adjusted p-value < 0.05, minimum average tags > 4). We found a total of 14731 ATAC-seq peaks had significant changes in accessibility due to LPS injection in Kupffer cells from the two parental strains. We found that 8125 peaks were similarly LPS responsive in both strains while more than 6,500 peaks met the statistical cutoff in only one strain.

Because fold change is determined by the ratio of expression under LPS and control treatment conditions, strain specific variation in either or both values contribute to detection of strain specific differences. Representative examples of genes exhibiting strain-similar or strain-specific responses to LPS are illustrated in Fig. 6c. *Tnf* exhibits nearly identical fold changes to LPS injection (∼60-fold) in BALB/cJ and C57BL/6J Kupffer cells respectively. Although *Tnf* is categorized as a strain-similar gene, basal and induced levels of *Tnf* in BALB/cJ Kupffer cells are approximately half those observed in C57BL/6J Kupffer cells. Thus, from a functional perspective, responses to Kupffer cell derived *Tnf* would be expected to be greater in C57BL/6J mice than BALB/cJ mice.

Recent studies of the responses of genetically diverse bone marrow derived macrophages to IL-4 addressed the relationship of fold response to absolute expression by categorizing strain specific RNA and epigenetic responses according to ‘low basal’, ‘similar basal’ and ‘high basal’ states (Hoeksema *et al.*, 2021) Following this approach, we defined ‘low basal’ mRNAs based on significantly lower basal expression in the LPS non-responsive strain relative to the responsive strain. An example of this type of gene is provided by *Slc7a2*, which exhibits almost no expression under basal or LPS treatment conditions in BABL/cJ cells but has measurable basal expression and is strongly induced by LPS in C57BL/6J cells. The ‘similar basal’ category is exemplified by *Med21*. Here, there is statistically equivalent basal expression in Kupffer cells from each strain, but LPS induction only in BALB/cJ Kupffer cells. The ‘high basal’ category is represented by *Plac8* where basal expression was significantly higher in C57BL/6J cells compared to BALB/cJ cells, and LPS treatment increased expression only in BALB/cJ cells. This results in *Plac8* being designated as LPS responsive in BALB/cJ Kupffer cells alone, even though it is more highly expressed in C57BL/6J cells in the LPS treatment condition. ATAC-seq peaks for the *Slc7a2*, *Med21* and *Plac8* promoters and nearby enhancers are shown in Fig. 6d and illustrate corresponding low basal, similar basal and high basal normalized tag counts, respectively. Distributions of the normalized tag counts and absolute numbers for each ATAC-seq peak category are illustrated in Fig. 6e-f. ’Equal basal’ enhancers accounted for ∼75% of ATAC-seq peaks exhibiting strain specific LPS responses. ‘High basal’ enhancers were nearly ∼25% of these peaks and ‘low basal’ peaks represented 4% (Fig. 6f).

Application of MAGGIE to these basal status peak categories revealed a clear segregation of motif mutations. Deleterious motif mutations for signal-dependent transcription factors, including IRFs, JUN/JUNB, and NFκB were enriched in equal basal ATAC peaks from the non-responsive strain (Fig. 6f, lower panel). These factors have established transcriptional roles in LPS-mediated TLR4 signaling (Glass and Natoli, 2016; Akira, Uematsu and Takeuchi, 2006; Kawai and Akira, 2008). In addition, MAFG:NFE2L1 were enriched suggesting previously unrecognized roles in the Kupffer cell response to LPS. Conversely, motif mutations improving motif scores in ‘high basal’ peaks from the non-responsive strain were enriched for lineage-determining transcription factors, including PU.1, ETS and C/EBP family members (Fig. 6f). A DR4 element was the second most significant motif mutation identified in ‘high basal’ ATAC peaks. Among nuclear receptors that recognize this motif, LXRα is the most highly expressed in Kupffer cells and is an established Kupffer cell lineage determining transcription factor (Lavin *et al.*, 2014; Mass *et al.*, 2016; Sakai *et al.*, 2019). The enrichment of mutations that improve the motif scores for Kupffer cell lineage determining factors in the high basal ATAC peaks is consistent with enhanced binding of these factors under basal conditions. A DR1 element recognized by RXR homodimers and heterodimers was enriched in ‘low basal’ ATAC-seq peaks and suggests a function in selection of a small subset of LPS-responsive enhancers. This result should be interpreted cautiously due to the limited number of peaks in this category.

The motif mutation analyses here do not discern whether genetic variants are acting in *cis* or *tra*ns. To estimate relative contributions of cis and trans effects in LPS responses, we performed ATAC-seq on Kupffer cells from CB6F1/J hybrids before and after LPS injection. We intersected peaks with *cis*-, *trans*-, or *mixed*-regulation from LPS treated with each category of low basal, similar basal and high basal peaks. The requirement to discern parental-specific alleles reduced the number of ATAC-peaks exhibiting strain-specific responses to 459 of 1261 strain-specific peaks. Of these, 133 peaks exhibited *cis* regulation, 40 *trans* regulation, and 22 *mixed* regulation (Pearson Chi^2^ p-value < < 3.00e-24, Fig. 6g). These observations suggest that strain-specific LPS responsiveness primarily results from local motif mutations in binding sites for signal dependent transcription factors.

**Fig. 6.**
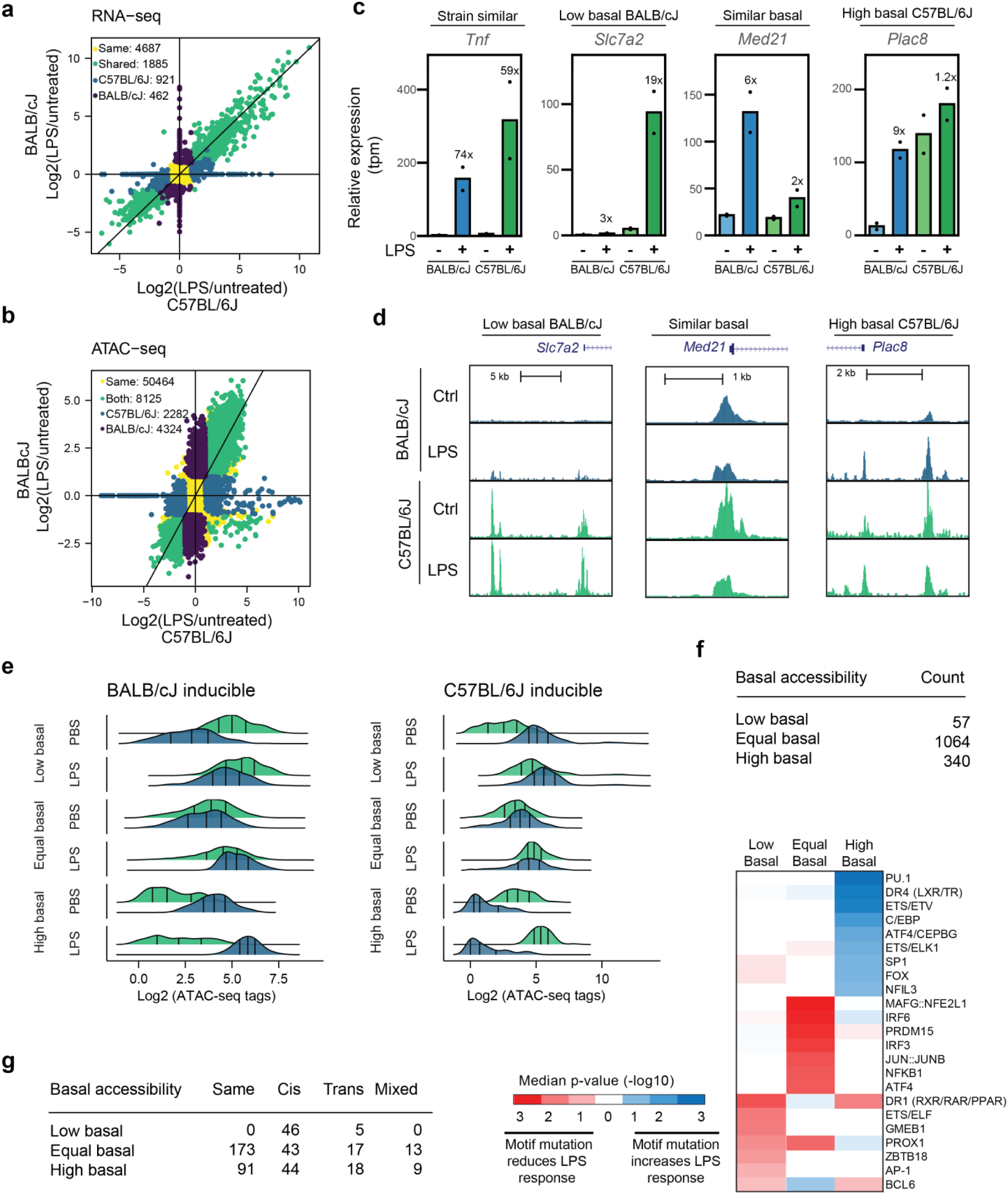
Strain-specific response to lipopolysaccharide determined by *cis* acting changes in motif affinity. **a**, Comparison of transcriptional response to LPS in C57BL/6J (x axis) and BALB/cJ (y axis) Kupffer cells, mice were treated with 0.1 mg/kg of LPS by intraperitoneal injection for 2 hours prior to isolation. Genes were considered differentially expressed if their expression differed with an absolute Log2 Fold Change > 1 and FDR adjusted p-value < 0.05. Genes were separated into C57BL/6J or BALB/cJ specific LPS-induced genes (blue and purple, respectively) or “same” if LPS induced in both C57BL/6J and BALB/cJ kupffer cells. **b**, comparison of ATAC-seq response to LPS in C57BL/6J (x-axis) and BALB/cJ (y-axis) Kupffer cells. **c**, Examples of strain similar and strain-differential transcriptional responses to LPS. **d**, ATAC-seq signal at nearby enhancers associated with low basal, equal basal, and high basal genes. **e,** Distribution of ATAC-seq signal in low basal, equal basal, and high basal enhancers. **f**, number of enhancers in each category and MAGGIE motifs associated with each category of enhancer. **g**, overlap of three enhancer categories with *cis* and *trans* enhancer analysis performed in CB6F1/J Kupffer cells isolated from mice treated with LPS for two hours. Pearson Chi^2^ p value for association of basal states and *cis*/*trans* regulation < 3.00e-24.

## DISCUSSION

Here, we have characterized the impact of natural genetic variation on gene expression and transcriptional regulatory elements in Kupffer cells derived from three widely used inbred strains of mice that exhibit different sensitivities to diet induced liver disease. The degree of variation between these strains is comparable to that between a given human individual and the human reference genome (Auton *et al.*, 2015), suggesting that the magnitude of effects on gene expression observed herein will be comparable to effects of common forms of genetic variation in human Kupffer cells. The observation that strain-specific differences in Kupffer cell gene expression were different from strain specific differences in primary cultured macrophages reinforces the concept that interpretation of common genetic variants associated with traits and disease risk requires characterization of cell specific regulatory landscapes, even for related cell types.

By acquiring and analyzing transcriptomic data for hepatocytes, stellate cells and endothelial cells in each strain and considering candidate hormonal signaling molecules, we inferred ligand receptor pairs predicted to contribute to strain-specific differences in Kupffer cells. Among these, we validated the prediction of enhanced leptin signaling in livers of BALB/cJ mice based on the preferential expression of *Lepr* in BALB/cJ Kupffer cells. It will be of interest to further investigate the physiologic significance of leptin signaling in Kupffer cells given a report that Kupffer cells facilitate the acute effects of leptin on hepatic lipid metabolism (Metlakunta *et al.*, 2017). Intriguingly, *Lepr* is one of the few genes that do not become expressed in monocyte derived cells that engraft the liver following loss of embryonic Kupffer cells, raising the questions of whether its expression requires embryonic origin and whether its function in Kupffer cells is limited to early in life prior to replacement with HSC-derived Kupffer cells.

Studies of NSG transplant and F1-hybrid models provided evidence that most strain specific differences in gene expression, open chromatin, and histone H3K27ac resulted from trans effects of genetic variation, and that the majority of these effects were a consequence of extracellular factors. A major objective of these studies was to investigate whether strain-specific differences in enhancers could be used to infer mechanisms underlying trans effects of genetic variation. From the analysis of ATAC-seq and H3K27ac ChIP-seq peaks exhibiting criteria for *trans* regulation in F1-hybrids, we detected clear biases for AP-1 and MAF family activation in BALB/cJ Kupffer cells and IRF, NFκB and LXR activity in C57BL/6J Kupffer cells. With the exception of increased expression of Maf in BALB/cJ Kupffer cells and LXRα in C57BL/6J Kupffer cells, these differential motif enrichments cannot be explained by differences in mRNA levels for the corresponding factors. Therefore, the strain specific differences in *trans*-regulated enhancer landscapes are most consistent with differences in extracellular environmental signals and intracellular signaling pathways.

Although we found a dominant role of *trans*-regulation in determining basal states of strain-specific gene expression and transcriptional regulatory elements, it was possible to exploit *cis* effects to establish the functional significance of motifs required for enhancer selection and function, as previously documented in BMDMs (Link *et al.*, 2018a). In addition, we found that *cis*-regulation predominated in the strain-specific responses to LPS. By segregating strain-specific responses according to the effects of genetic variation on relative activity under basal conditions, we identified qualitatively different patterns of motif mutations associated with each category. Mutations disrupting binding sites for transcription factors mediating transcriptional responses to LPS were significantly associated with impaired activation, as expected. In addition, mutations improving suboptimal binding sites for Kupffer cell lineage determining factors resulted in high constitutive basal activity and a reduced dynamic range in response to LPS. This observation is similar to recent findings in IL-4 treated BMDMs (Hoeksema *et al.*, 2021) and suggests the importance of suboptimized binding sites for lineage determining factors in conferring low levels of basal activity but enabling the selection of primed enhancers that are receptive to binding and activation by signal dependent transcription factors.

In concert, systematic analysis of the transcriptomes and regulatory landscapes of Kupffer cells in A/J, BALB/cJ and C57BL/6J mice, in NSG chimeras, and in F1-hybrids, reveals dominant *trans* effects of natural genetic variation under homeostatic conditions and dominant *cis* effects on the response to LPS. The transcriptomic and genomic data sets resulting from these studies provide valuable resources for further understanding the mechanisms by which genetic variation affects tissue resident macrophage phenotypes and susceptibility to diseases in which macrophages play pathogenic or protective roles.

## ACKNOWLEDGEMENTS

These studies were supported by NIH grants DK091183 and HL147835 and a Leducq Transatlantic Network grant 16CVD01 to CKG. Sequencing costs were partially supported by DK063491. TDT was supported by P30 DK063491, T32DK007044, and NRSA T32CA009523. HB was supported by supported by the NIH Predoctoral training grant T32DK007202 and F30DK124980. JSS was supported by American Heart Association Fellowship 16PRE30980030 and NIH Predoctoral Training Grant 5T32DK007541. M.S. was supported by the Manpei Suzuki Diabetes Foundation of Tokyo, Japan, and the Osamu Hayaishi Memorial Scholarship for Study Abroad, Japan. This study was also supported by NIH Grant DK120515.

## AUTHOR CONTRIBUTIONS

Conceptualization: T.D.T, J.S.S., M.S., H.B., and C.K.G. Formal analysis T.D.T, H.B., J.S.S., E.Z., H.B., and C.K.G.

Investigation: T.D.T., H.B., E.Z., M.S., J.S.S., and Y.A. Writing: T.D.T., H.B., E.Z., and C.K.G. Visualization: H.R.B. and T.D.T. Supervision: C.K.G. and B.S. Funding Acquisition: C.K.G.

## RESOURCE AVAILABILITY

All data needed to evaluate the conclusions in the paper are present in the paper. Sequencing data will be published on the gene expression omnibus (GEO) at time of publication. Additional data related to this paper may be requested from the author.

### Lead Contact

Further information and request for resource and reagents should be directed to and will be fulfilled by the Lead Contact, Christopher K. Glass

### Materials Availability

This study did not generate new unique reagents.

## Methods

### Mice

A/J, BALB/cJ, C57BL/6J, and CB6F1/J mice used in this study were sourced directly from Jackson laboratories, with the exception of C57BL/6 (lab maintained) mice used in NASH experiments. Immunodeficient NOD-*scid* IL2Rg^null^ mice were obtained from the UC San Diego Moore’s Cancer Center. All animals were maintained and all procedures performed in accordance with an approval animal study protocol meeting AALAC standards.

### Individual Bone Marrow Chimeras

NOD-scid IL2Rgnull mice were conditioned with the myeloablative agent busulfan at 25 mg/kg for 2 consecutive days, as described (Montecino-Rodriguez and Dorshkind, 2020). On the third day, mice were engrafted via retro orbital injection with magnetically enriched, lineage negative, hematopoietic stem cells (Miltenyi) from BALB/cJ or C57BL/6J donors. Engraftment efficiency was monitored in peripheral blood after 4 and 8 weeks. Livers from chimeric recipients were harvested 4 months after transplant, and graft derived Kupffer cells purified by fluorescence activated cell sorting (FACS). In BALB/cJ chimeras, viable graft derived Kupffer cells were distinguished by MHC haplotype as CD146-, F4/80+, Cd11b+, H2-Db-(KH95), H2-Dd+(34-2-12). In C57BL/6J chimeras, graft derived cells were distinguished as H2-Kd-(SF1-1.1), H2-Kb+(AF6-88.5). Kupffer cells from each group (n=3) were used downstream for RNA-seq.

### NASH-Model Diets

Mice were fed for up to 30 weeks with a NASH-model diet (Research Diets, D09100301) composed of 40 kcal% from fat, 20 kcal% from fructose, and 2% cholesterol by mass, or a custom defined control diet (Research Diet, D15100601) composed of 10% kcal from fat with 50 g inulin (a dietary fiber) per 4,057 kcal.

### Histology and Pathologic Scoring

Samples from NASH and control mouse livers were incubated at room temperature in 1% PFA for 24 hours. Sections were paraffin embedded and sectioned by the UCSD Histology Core. Sections were stained with hematoxylin and eosin or picosirius red to evaluate for steatosis and fibrosis, respectively. Samples were scored by a board certified pathologist blinded to the sample group using the NASH CRN scoring system and fibrosis scoring system (Kleiner *et al.*, 2005).

### *In vivo* response to leptin

Leptin stocks were prepared in Tris-HCl pH8.0 buffer and diluted to 0.333mg/mL in PBS for intraperitoneal injection at 1 mg/kg into overnight fasted (18:00 to 6:00). At the indicated times, mice were humanely euthanized by CO_2_, and liver tissues removed, minced, then homogenized in lysis buffer (50 mM HEPES-KOH (pH 7.9), 150 mM NaCl, 1.5 mM MgCl_2_, 1% NP-40, 1mM phenylmethylsulfonyl fluoride (PMSF) (Sigma-Aldrich), protease inhibitor cocktail (Sigma-Aldrich), PhosSTOP (Roch)). Lysates were sonicated by ultrasound homogenizer (Bioruptor, Diagenode) for 10 min at 4°C, and centrifuged for 10 min at 15000 rpm at 4°C. The supernatant was used as tissue homogenate for immunoblotting. After a protein assay (Bio-Rad Laboratories), the homogenate was boiled at 95°C for 5 min in NuPAGE^TM^ LDS Sample Buffer (Thermo Fisher Scientific) with NuPAGE^TM^ Sample Reducing Agent (Thermo Fisher Scientific), subjected to SDS-PAGE, and transferred to immobilon-P transfer membranes (Merck Millipore). Immunodetection was carried out with anti-phospho-STAT3 (Tyr705) (Cell Signaling Technology, #9145), anti-STAT3 (Cell Signaling Technology, #9139) or anti-b-actin (Sigma-Aldrich, A2228) and bound antibodies were visualized with peroxidase-conjugated affinity-purified donkey anti-mouse or anti-rabbit IgG (Dako) using Luminate^TM^ Forte Western HRP Substrate (Merck Millipore), and luminescence images were analyzed by ChemiDoc XRS+ System (Bio-Rad Laboratories).

### *In vivo* response to acute lipopolysaccharide treatment

Male mice were fasted overnight and treated for 2 hours by intraperitoneal injection of 0.1 mg/kg *E. coli* O114:B4 lipopolysaccharide. Control mice were equivalently fasted but uninjected. Mice were euthanized by CO_2_ exposure and transcription was halted by hepatic perfusion with flavopiridol (1 mg/ml). Liver tissue was digested in situ with Liberase TM in the presence of flavopiridol, and immunolabeled Kupffer cells were purified by fluorescence activated cell sorting (Sakai *et al.*, 2019; Seidman *et al.*, 2020).

### Hepatocyte Preparation

Hepatocytes were prepared by perfusion digestion in a retrograde fashion through the inferior vena cava to the portal vein. In brief, livers were blanched with clearing buffer (HBSS + 10 mM HEPES), then digested with a collagenase solution (HBSS supplemented with 0.3 mg/ml collagenase D, 10 mM HEPES, and 1 tablet protease inhibitor cocktail complete-EDTA free per 50 ml) at 39C. Livers were perfused for 18 minutes at 5 ml/min. The perfusion and digestion steps were performed in the presence of 1 mM flavopiridol to offset transcriptional changes associated with digestion. After digestion, individual livers were gently dissociated using forceps in 20 ml Medium 199 supplemented with 5% FBS and penicillin/streptomycin/gentamycin. Crude hepatocyte preps were carefully strained through a 100-micron strainer into a 50 ml tube. An equal volume of isotonic Percoll (90% Percoll, 10% 10X HBSS) was added, followed by gentle mixing, then centrifugation at 100xG for 7 min at 4°C. The supernatant was discarded, and the cells were gently resuspended with 50 ml Medium 199 and centrifuged at 100xG for 2 min at 4C. Hepatocytes were gently resuspended in Medium 199 and counted.

### Hepatic non-parenchymal cell preparation

Non-parenchymal cells from digested livers were prepared as described (Sakai *et al.*, 2019; Seidman *et al.*, 2020; Troutman *et al.*, 2021). In brief, livers were retrograde perfused for 3 min at a rate of 5-7 mL/min through the inferior vena cava with HBSS without Ca2^+^ or Mg2^+^ supplemented with 0.5 mM EGTA, 0.5 mM EDTA, and 20 mM HEPES. Perfusions were then switched to 40 mL of a digestion buffer, held at 37°C, comprised of HBSS with Ca2^+^ and Mg2^+^ supplemented with 0.033 mg/mL of Liberase TM (Roche), 20 mg/mL DNaseI (Worthington), and 20 mM HEPES. Livers were then excised, minced, and digested for an additional 20 min in vitro at 37°C with gentle rotation in 20 mL of fresh digestion buffer. The perfusion and digestion steps were performed in the presence of 1 mM flavopiridol to offset transcriptional changes associated with digestion. After tissue digestion, cells were passed through a 70-micron cell strainer and hepatocytes removed by 2 low-speed centrifugation steps at 50 X G for 2 min. Non-parenchymal cells in the supernatant were further separated from debris by pelleting for 15 min at 600 X G in 50 mL of 20% isotonic Percoll (Sigma Aldrich) at room temperature. Cells were then washed from Percoll containing buffer and suspended in 10 mL 28% OptiPrep (Sigma Aldrich) and carefully underlaid beneath 3 mL of wash buffer. The resulting gradient was centrifuged at 1,400 X G for 25 min at 4°C with no break, and cells enriched at the interface were saved and subjected to isotonic erythrocyte lysis. Cells were washed after erythrocyte lysis and immediately used purified by cells sorting.

### Cell Sorting and Flow Cytometry

Hepatic non-parenchymal cells were labeled with fluorescent antibodies and desired cell populations were purified using a Beckman Coulter Mo-Flo Astrios EQ configured with spatially separated 355 nm, 405 nm, 488 nm, 561 nm, and 642 nm lasers. Kupffer cells were defined as 355:494/20Low, SSCLow, CD146Neg, CD45Pos, F4/80High, CD11bIntermediate, Live, Singlets. Liver sinusoidal endothelial cells were defined at 355:494/20Low, SSCLow, CD45Neg, CD146Pos, Live, Singlets. Hepatic stellate cells were defined as 355:494/20High, SSCIntermediate, Live, Singlets.

### Bone Marrow Derived Macrophages

Femur, tibia and iliac bones from mice were flushed with DPBS, and red blood cells were lysed using red blood cell lysis buffer (Sigma Aldrich). Bone marrow cells were seeded in 10 cm non-tissue culture plates in RPMB with 10% FBS, 30% L929-cell conditioned laboratory-made media (as source of M-CSF), 100 U/ml penicillin-streptomycin (Thermo Fisher Scientific), and 16.7 ng/ml M-CSF (BioLegend). After 2-3 days of differentiation, cells were fed with 16.7 ng/ml M-CSF. After an additional 2 days of culture, non-adherent cells were washed off with room temperature PBS, and adherent macrophages were obtained by scraping. Cells were counted, density adjusted with RPMI supplemented with 10% FBS, 100 U/ml penicillin-streptomycin, seeded into multiwell plates, and rested at 37°C overnight. The following day, macrophages were treated with 100 ng/ml Kdo_2_-lipid A (Avanti lipids), a highly purified E. coli lipopolysaccharide (Raetz *et al.*, 2006).

### Next Generation Sequencing Libraries

#### ATAC-seq

Transposase reactions and sequencing libraries were generated as described previously (Buenrostro et al., 2013; Sakai et al., 2019; Seidman et al., 2020) using 25,000 to 50,000 FACS purified Kupffer cells. Tagmented DNA was cleaned up using Zymo ChIP Clean & Concentrate columns and PCR amplified for 14 cycles using barcoding primers. Libraries were size selected to 175-225 bp using gel excision and purified as described (Texari *et al.*, 2021). For F1 samples, dual indexed libraries were pooled for a targeted depth of 100 million reads per sample.

#### ChIP-seq

Chromatin immunoprecipitation and sequencing libraries were generated as previously described (Texari *et al.*, 2021) with the modifications to lysis, immunoprecipitation buffer, and washing buffer as described (Eichenfield *et al.*, 2016; Seidman *et al.*, 2020). In brief, FACS purified cells were fixed with 1% paraformaldehyde for 10 min at room temperature. Next, 2.625Mglycine was added to 125mM to quench fixation and cells were collected by centrifugation with the addition of 0.01% Tween-20 at 1,200 X G for 10 min at 4°C. Cells were washed once with 0.01% Tween-20 in PBS and collected by centrifugation at 1,200 XG for 10 min at 4°C. Cell pellets were then snap frozen and stored at -80C. For ChIP reactions, cell pellets were thawed on ice and lysed in 80 ml LB3 (10mMTris/HCl pH 7.5, 100mMNaCl, 1mMEDTA, 0.5mM EGTA, 0.1% deoxycholate, 0.5% sarkosyl, 1 x protease inhibitor cocktail, and 1 mM sodium butyrate). Lysate was sonicated using a Covaris for 12 cycles with the following setting: time, 60 s; duty, 5.0; PIP, 140; cycles, 200; amplitude, 0.0; velocity, 0.0; dwell, 0.0. Samples were collected and 10% Triton X-100 was added to 1% final concentration. One percent of the sonicated lysate was saved as a ChIP input. For each chromatin immunoprecipitation, aliquots of ∼500,000 cells were added to 20 µl Dynabeads Protein A with 2 µg anti-H3K27ac (Active Motif) and incubated with slow rotation at 4°C overnight. The following day, beads were collected using a magnet and washed three times each with wash buffer I (20 mM Tris/HCl pH 7.5, 150 mM NaCl, 1% Triton X-100, 0.1% SDS, 2 mM EDTA, and 1 x protease inhibitor cocktail) and wash buffer III (10 mM Tris/HCl pH 7.5, 250 mM LiCl, 1% Triton X-100, 0.7% Deoxycholate, 1 mM EDTA, and 1 x protease inhibitor cocktail). Beads were then washed twice with ice cold 10 mM Tris/HCl pH 7.5, 1 mM EDTA, 0.2% Tween-20. Sequencing libraries were prepared for ChIP products while bound to the Dynabeads Protein A as in (Texari *et al.*, 2021). For F1 samples, dual indexed libraries were pooled for a targeted depth of 100 million reads per sample.

#### RNA-seq

Poly A RNA-seq libraries were generated as previously described (Gosselin *et al.*, 2014; Oishi *et al.*, 2016) using 50,00 to 100,000 FACS purified cells stored in lysis/Oligo d(T) Magnetic Beads binding buffer and stored at -80°C, or 500 ng of purified RNA using the Zymo Research Direct-zol RNA microprep kit. In brief, mRNAs were enriched by incubation with Oligo d(T) Magnetic Beads (NEB, S1419S) and then fragmented/eluted by incubation at 94°C for 9 min. Poly A enriched mRNA was fragmented, in 2x Superscript III first-strand buffer with 10 mM DTT (Invitrogen), by incubation at 94°C for 9 min, then immediately chilled on ice before the next step. The 10 µL of fragmented mRNA, 0.5 µL of random primer (Invitrogen), 0.5 µL of Oligo dT primer (Invitrogen), 0.5 µL of SUPERase-In (Ambion), 1 µL of dNTPs (10 mM) and 1 µL of DTT (10 mM) were heated at 50°C for 3 minutes. At the end of incubation, 5.8 µL of water, 1 µL of DTT (100 mM), 0.1 µL Actinomycin D (2 mg/mL), 0.2 µL of 1% Tween-20 (Sigma) and 0.2 µL of Superscript III (Invitrogen) were added and incubated in a PCR machine using the following conditions: 25°C for 10 min, 50°C for 50 min, and a 4°C hold. The product was then purified with RNAClean XP beads according to manufacturer’s instruction and eluted with 10 µL nuclease-free water. The RNA/cDNA double-stranded hybrid was then added to 1.5 µL of Blue Buffer (Enzymatics), 1.1 µL of dUTP mix (10mMdATP, dCTP, dGTP and 20 mM dUTP), 0.2 µL of RNase H (5 U/mL), 1.05 µL of water, 1 µL of DNA polymerase I (Enzymatics) and 0.15 µL of 1% Tween-20. The mixture was incubated at 16°C for 1 h. The resulting dUTP-marked dsDNA was purified using 28 µL of Sera-Mag Speedbeads (Thermo Fisher Scientific), diluted with 20% PEG8000, 2.5M NaCl to final of 13% PEG, eluted with 40 µL EB buffer (10 mM Tris-Cl, pH 8.5) and frozen at -80°C. The purified dsDNA (40 µL) underwent end repair by blunting, A-tailing and adaptor ligation as previously described (Heinz *et al.*, 2010) using indexed barcoding adapters. Libraries were PCR-amplified for 9-14 cycles, size selected by gel extraction, quantified using a Qubit dsDNA HS Assay Kit (Thermo Fisher Scientific) and sequenced on a Hi-seq 4000, NextSeq 500, or a NOVA-seq (Illumina, San Diego, CA) according to the manufacturer’s instructions. For F1 samples, dual indexed libraries were pooled for a targeted depth of 100 million reads per sample.

## QUANTIFICATION AND STATISTICAL ANALAYSIS

### Sequencing Data Analysis

#### Preprocessing and Mapping

Sequencing data were assessed for quality using fastqc. ATAC-seq and ChIP-seq data were mapped using Bowtie2 and RNA-seq data was mapped using STAR (Langmead and Salzberg, 2012; Dobin *et al.*, 2013). ATAC-seq data were trimmed to 30 bp to remove sequencing adapters, which improved mapping efficiency. Strain specific genomes for BALB/cJ and A/J were generated from by replacing invariant positions of mm10 sequence with alleles reported in the Mouse Genome Project strain specific VCF files. mm10 was used as the C57BL/6J strain specific genome. Samples from parental strains of mice were mapped to the strain specific genome. Mapped reads were shifted to the chromosome coordinates of the mm10 genome build using MARGE.pl shift with -ind set to balbcj or aj for reads mapped to the BALB/cJ or A/J genome, respectively (Link *et al.*, 2018b).

For samples from CB6F1/J samples, reads were mapped to the mm10 and BALB/cJ genome builds. Then the BALB/cJ mapped reads were shifted to the mm10 build with MMARGE as above. Perfectly mapped reads spanning genetic mutations between BALB/cJ and mm10 were identified using the MMARGE.pl allele_specific_reads command with -ind set to BALB/cJ and a second time with -ind set to mm10 resulting in two SAM files for each biological sample: one SAM file containing reads perfectly mapped to the mm10 genome that spanned known DNA sequence differences relative to the BALB/cJ genome; and a second SAM file containing reads perfectly mapped to the BALB/cJ genome that spanning known DNA sequence differences relative to the reference mm10 genome.

#### ATAC-seq analyses

Strain specific ATAC-seq SAM files were used to generate HOMER tag directories and allelic IDR peaks were identified using each biological replicate. Alterations in allelic signals from pooled IDR peaks were detected using DeSeq2 and required the following thresholds: minimum normalized average tag depth > 16; absolute log_2_ fold-change > 1; and adjusted p-value < 0.05.

#### ChIP-seq analyses

In the F0 mice, H3K27Ac ChIP-seq tags were quantified for differential peak analysis by annotating merged ATAC-seq peaks with ChIP-seq tag directories using the HOMER command “annotatePeaks.pl” parameters -size 1000 -raw (Heinz *et al.*, 2010). H3K27Ac tag counts were quantified for visualization in heatmaps using “annotatePeaks.pl” with the following parameters -size 1000 -norm 1e7 (Heinz *et al.*, 2010). In F1 mice, strain specific SAM files were used to generate strain specific H3K27Ac tag directories that only contained perfectly aligned reads spanning a mutation between the intercrossed strains. H3K27Ac reads spanning mutations were aggregated over IDR peaks using a 2000bp window centered on the IDR peak. Alterations in allelic tag counts from pooled peaks were detected using DESeq2 with the following thresholds normalized average tag depth > 16; absolute log_2_ fold-change > 1; and adjusted p-value < 0.05.

#### RNA-seq analyses

Gene expression data was quantified using the HOMER command analyzeRepeats. Raw count data was aggregated using the following parameters: rna mm10 -condenseGenes -count exons -noadj. TPM count data was aggregated using the following parameters: rna mm10 -count exons -tpm (Heinz *et al.*, 2010). TPM values were matched to the isoforms with the highest raw count values. Only genes with an average expression level > 8 TPM were considered for differential gene analysis. Differentially expressed genes were identified using DESeq2 with betaPrior set to TRUE (Love, Huber and Anders, 2014). *Trans* gene ontology analysis was performed using HOMER findGO.pl (Heinz *et al.*, 2010). All other gene ontology enrichment analyses were performed using Metascape (Zhou *et al.*, 2019).

#### Motif enrichment analysis

Enrichment of known transcription factor binding motifs in ATAC-seq peaks was performed using HOMER. DNA sequences associated with peaks containing no detectable difference in LPS responsiveness in both BALB/cJ and C57BL/6J were used as the background for enrichment analysis of IDR ATAC-seq peaks from F0 Kupffer cells. A randomly generated GC-matched background was used for enrichment analysis of allele-specific IDR ATAC-seq peaks from F1 Kupffer cells. Motifs were selected for visualization if the probability of enrichment over background had a q-value < 0.05 in only one strain or allele.

#### Niche-net

NicheNet is a computational model that ranks ligands expressed by one or more cell types by their ability to induce target gene expression in a cell of interest (Browaeys, Saelens and Saeys, 2020). To assess putative strain specific ligand activity, we first filtered the NicheNet ligand-target matrix to only consider ligands in which:

1. The ligand was expressed by a cell of the hepatic niche within that strain at > 10 TPM.
2. The receptor was expressed by Kupffer cells from that strain at > 10 TPM.

We also included selected metabolic ligands for which expression data were not available. We did not require that ligands or their receptors be differentially expressed by sender or receiver cells. Target genes were selected to be any gene that had significantly higher expression in a pairwise comparison of that strain (“union” gene set, adjusted p value < 0.05, log fold change > 2, TPM > 10 expression in Kupffer cells). As a background we considered all genes that were expressed at TPM > 10 in Kupffer cells. The NicheNet ligand activity score was then computed as the Pearson correlation coefficient between the ligand-target score and the binary vector indicating whether a target gene was differentially expressed. For heatmaps two top scoring ligands from each strain were aggregated and displayed with ligand z-scores. Ligand receptor interaction scores were displayed for ligand-receptor pairs with a receptor expressed by Kupffer cells in at least one strain. For the circos plot analysis ligand-target interaction scores were displayed as arrow thicknesses linking a ligand to its target gene.

### Statistical Analysis

Genome wide signals for RNA-seq, ATAC-seq, and ChIP-seq were evaluated for differential levels using DESeq2 (Love, Huber and Anders, 2014). The raw p-values from DESeq2 for a given peak or gene were corrected for multiple testing using the Benjamini-Hochberg procedure. Effect of strain on NASH CRN and fibrosis score was assessed with a Kruskal-Wallis test (R, Kruskal.test) Association of weight change with strain, diet, and a composite variable of strain and diet was assessed with a linear mixed effects model (R, nmle). Association of *cis*/*trans* with low, equal, and high basal regulation was performed with a Pearson Chi^2^ test (Python scipy.stats.contingency.chi2_contingency).

#### Data visualization

Data were visualized using the UCSC genome browser (Kent *et al.*, 2002) and custom R and Python scripts.

### Data and Code Availability

All sequencing data will be made available within the Gene Expression Omnibus (GEO) database at time of publication. Code for analysis performed herein will be made available upon reasonable request.

## Extended Data

**Extended Data Figure 1.**
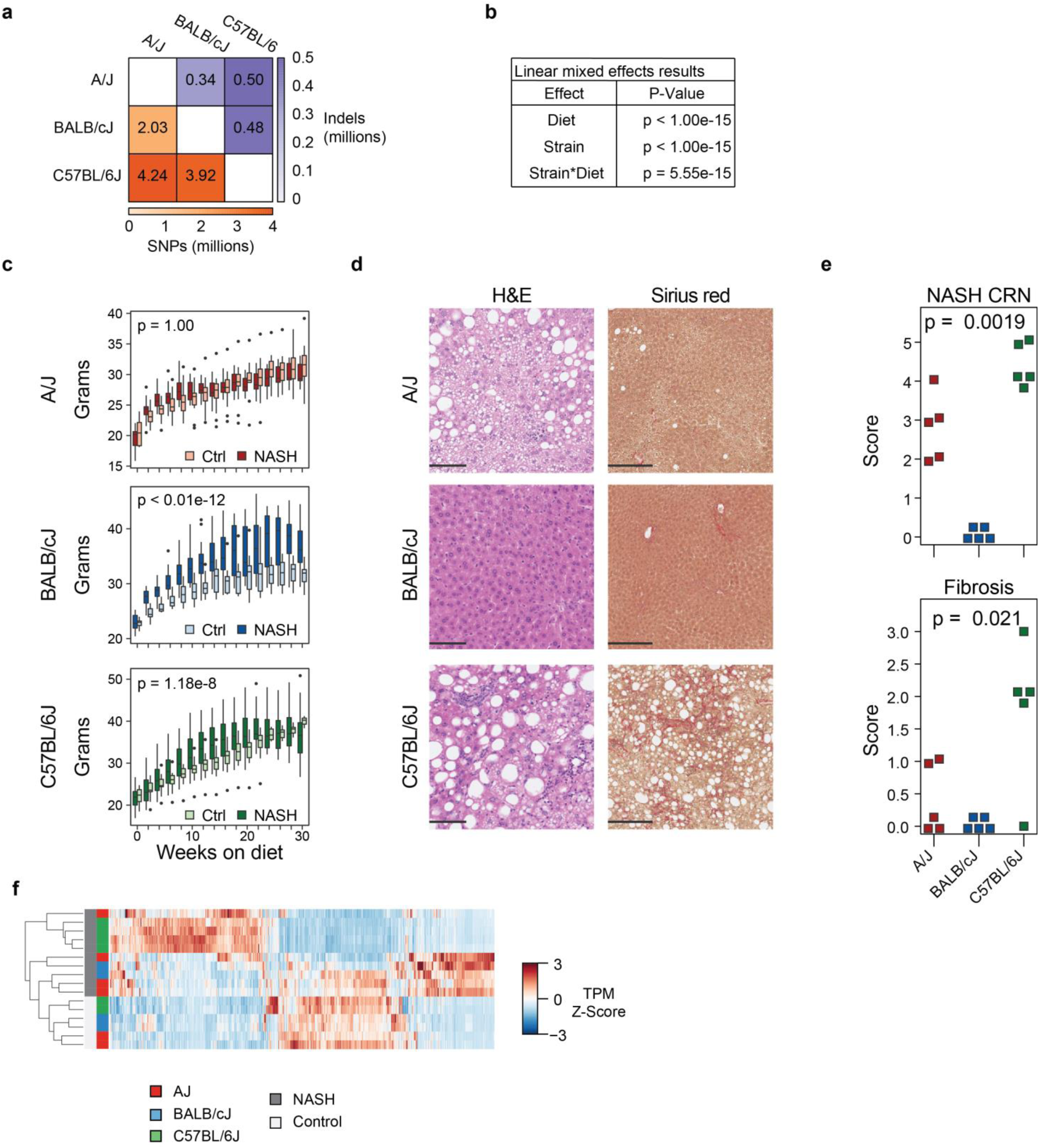
**a**, Degree of natural genetic variation between three inbred strains used in this study. **b**, Linear mixed effects model of effects of diet and genotype on mouse mass. **c**, Weekly weight gain in each strain on AMLN diet. **d**, Histopathological evidence of NASH in following 30 weeks of AMLN diet in each strain of mice using hematoxylin and eosin (left) and Sirius red (right) staining of mouse livers. **e**, Histopathological scoring of NASH (top) and fibrosis (bottom) in each strain of mouse following 30 weeks of AMLN diet. Strain effects were assessed independently for NASH CRN and fibrosis score using a Kruskal-Wallis test (R, kruskal.test). **f**, Unsupervised clustering of strain specific differential genes from bone marrow derived macrophages in three strains of mice or Kupffer cells. Differential genes were defined using pairwise comparisons with a Log2 Fold Change > 1 and an FDR adjusted p-value > 0.05.

**Extended Data Figure 2.**
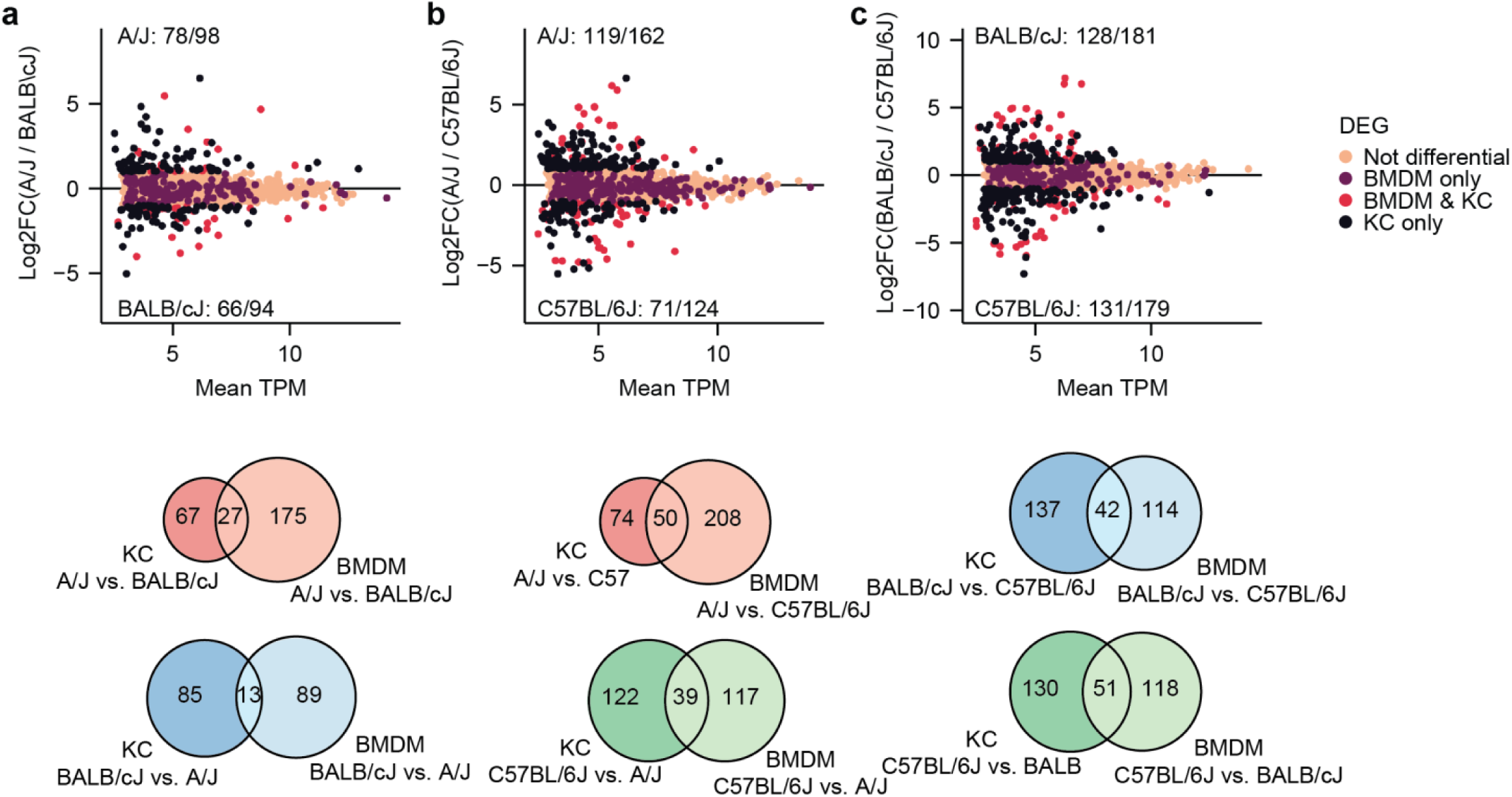
**a**, comparison of transcriptional effect of genetic variation in Kupffer cells and BMDMs from A/J and BALB/cJ mice. MA plot of differential gene expression between the strains was overlaid with information considering overlap with differential genes from comparison of A/J and BALB/cJ BMDMs (top). Pink dots indicate genes that are not differential in either strain, purple dots are only differential in BMDM, black dots are genes that are only differential in Kupffer cells, and red dots are genes that are differential in both BMDM and Kupffer cells (Bottom row). Total overlap of strain specific genes in Kupffer cells and BMDMs for each pairwise comparison of three inbred strains. **b**, plots for A/J and C57BL/6J comparison. **c**, plots for BALB/cJ and C57BL/6J comparison.

**Extended Data Figure 3.**
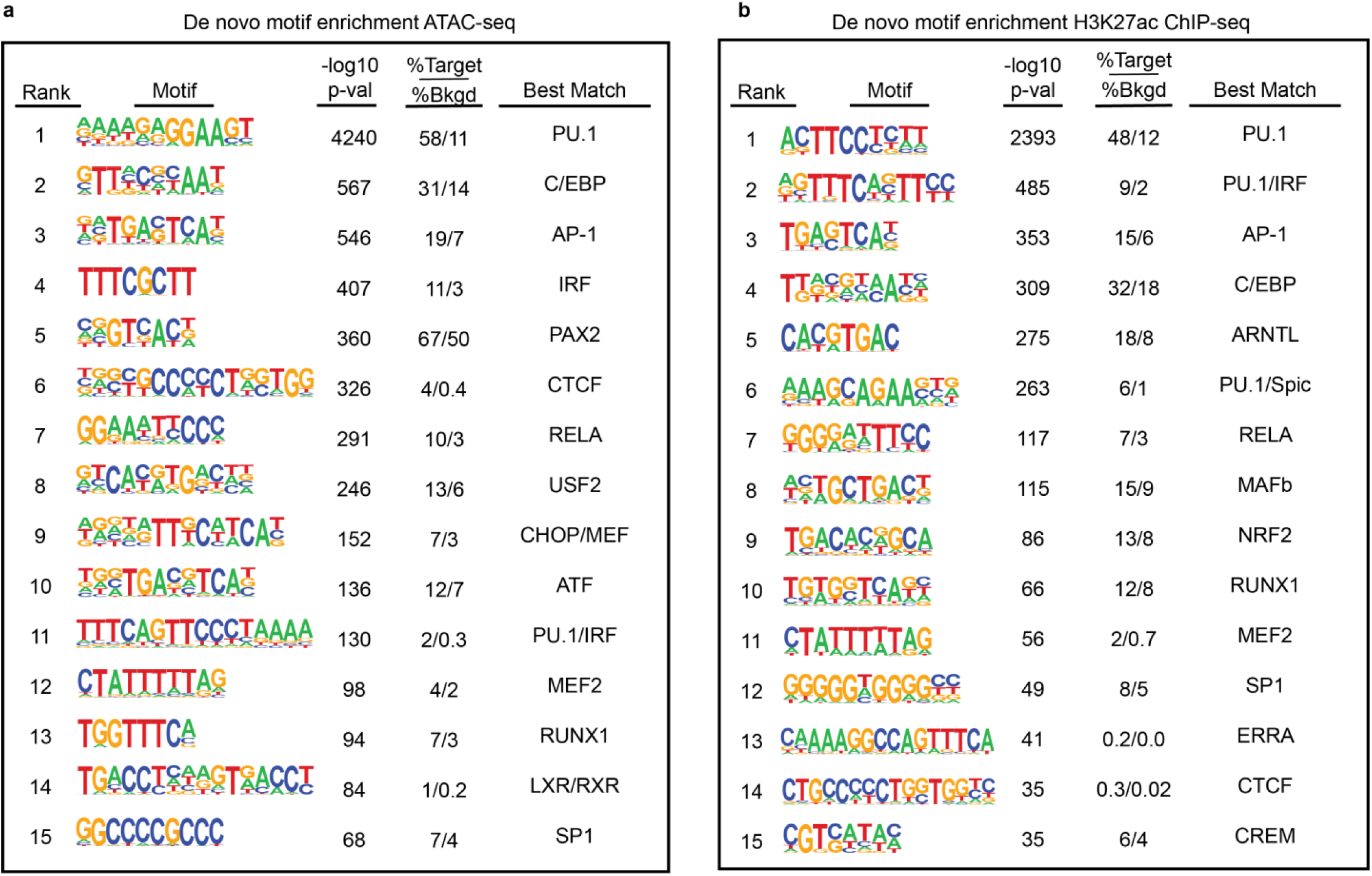
HOMER *de novo* motif analysis of shared poised enhancers (“ATAC”, ATAC-seq tags > 8) shared by all 3 strains and active enhancers (“H3K27Ac”, H3K27Ac ChIP-seq tags > 16). **a**, top 15 *de novo* motifs enriched at poised enhancers as determined by HOMER. **b**, top 15 *de novo* motifs enriched at poised enhancers as determined by HOMER.

**Extended Data Figure 4.**
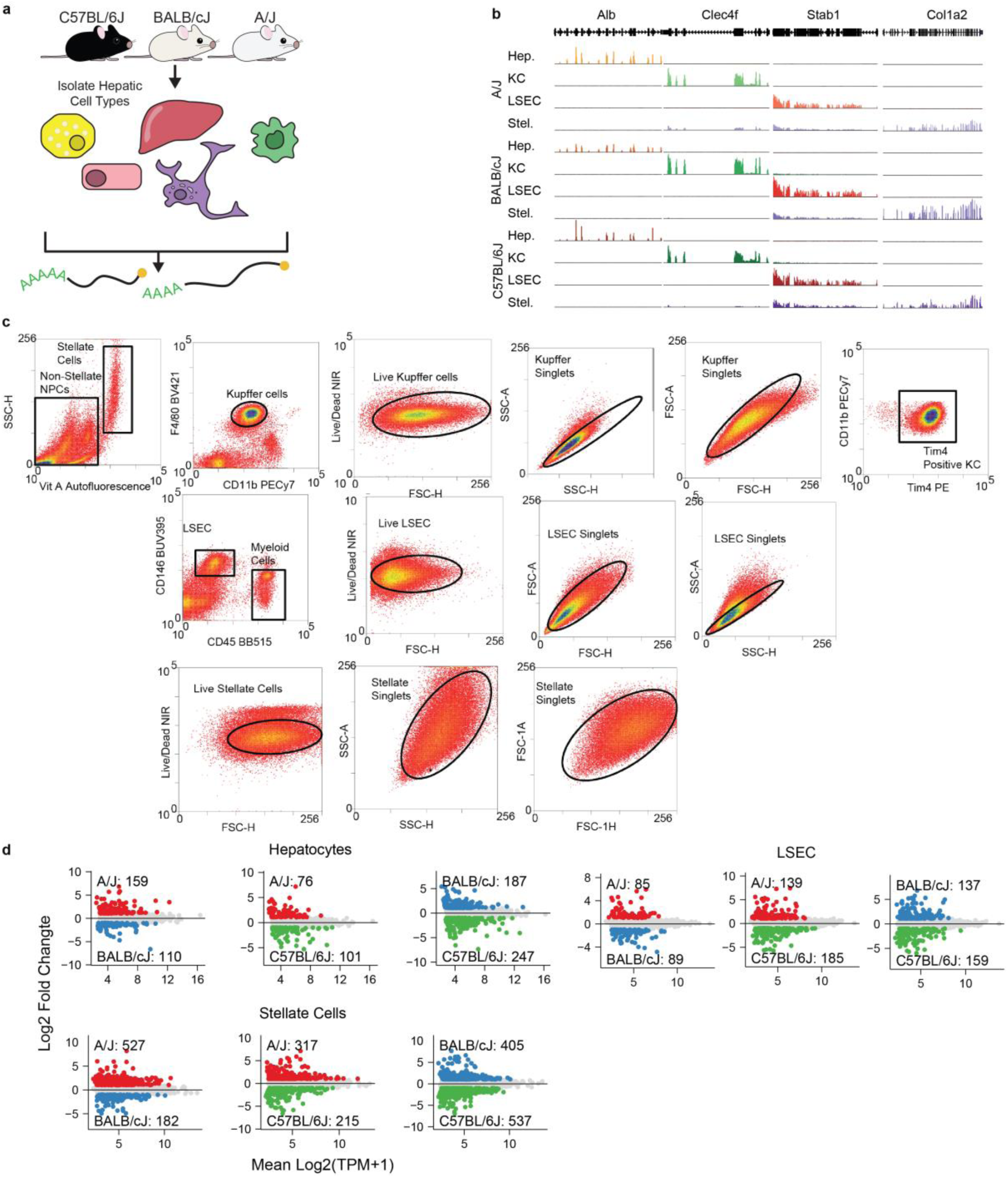
**a**, experimental schematic for isolation of hepatic cell types from inbred strains. **b**, assessment of cell isolation purity at cell specific gene expression loci. Hepatocytes (yellow), Kupffer cells (green), and LSECs (red) were sorted with <1% contamination. Hepatic stellate cell RNA-seq libraries displayed minor (<10%) contamination with LSECs and Kupffer cells (seen as RNA-seq signal in *Clec4f* and *Stab1* loci). **c**, FACS strategy for stellate, LSEC, and Kupffer cell isolation. **d**, strain-specific transcriptional variation in hepatocytes, LSECs, and Stellate cells.

**Extended Data Figure 5.**
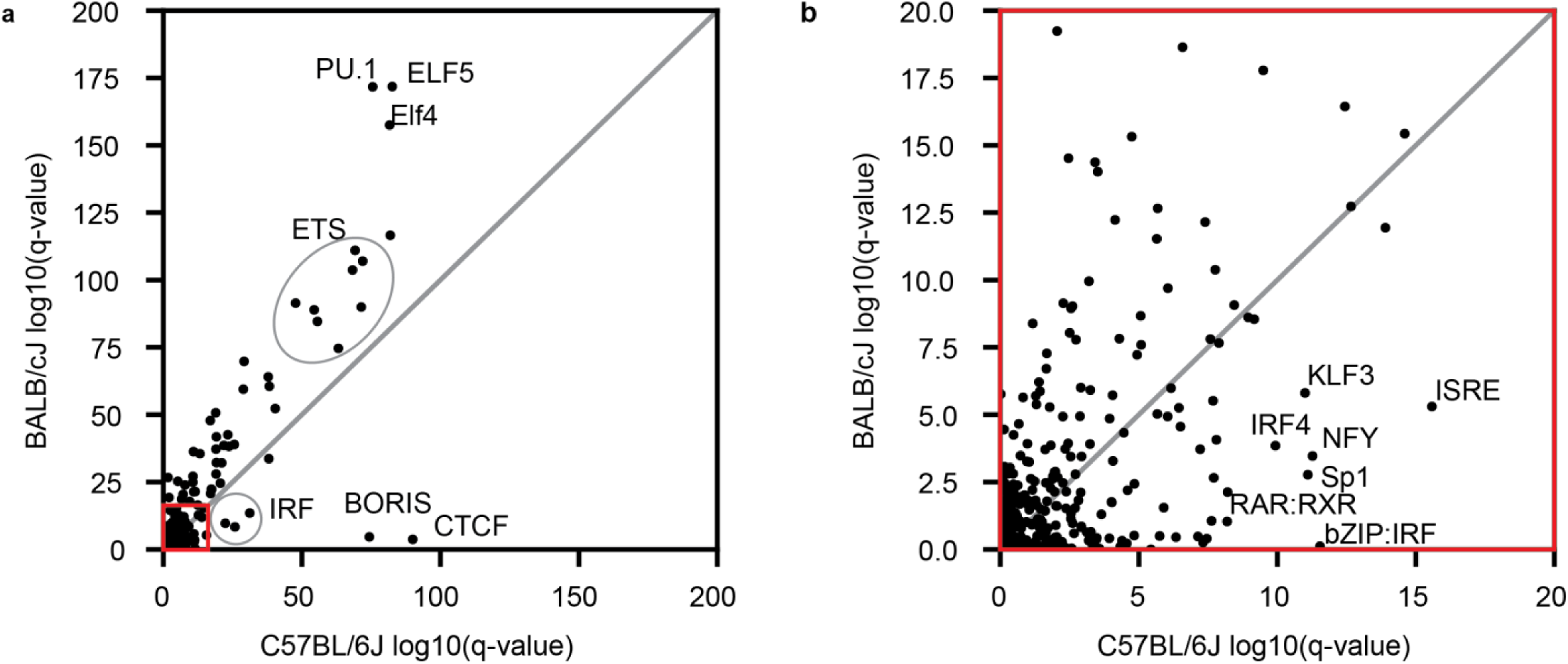
**a**, HOMER known motif enrichment in BALB/cJ *trans* regulated ATAC-seq peaks (y-axis) and C57BL/6J *trans* regulated ATAC-seq peaks (x-axis). **b**, view of highlighted region in a.

